# Fast calcium-dependent fluorescent labeling for recording of neuronal activation

**DOI:** 10.64898/2026.08.05.742984

**Authors:** Nicola Porzberg, Jennifer Heck, Jonas Wilhelm, Joergen Benjaminsen, Teresa Blümel, Magnus-Carsten Huppertz, Kyung-Min Noh, Thomas Thumberger, Martin Heine, Jochen Wittbrodt, Sinem Saka, Julien Hiblot, Kai Johnsson

**Affiliations:** Department of Chemical Biology, Max Planck Institute for Medical Research, Heidelberg, Germany; European Molecular Biology Laboratory, Heidelberg, Germany; Institute for Developmental Biology and Neurobiology, Johannes Gutenberg University, Mainz, Germany; Centre for Organismal Studies (COS), Heidelberg University, Heidelberg, Germany; Institute of Chemical Sciences and Engineering (ISIC), École Polytechnique Fédérale de Lausanne (EPFL), Switzerland

## Abstract

Calcium transients encode cellular and neuronal activity across timescales ranging from milliseconds to hours, yet linking these transient signals to downstream molecular states remains a major challenge. We recently introduced Caprola, a calcium-dependent protein labeling tool that converts calcium transients into permanent fluorescent marks for later analysis. In this way, Caprola enables tracking of neuronal activities in animal models as well as retrospective identification of labeled cells for isolation and transcriptomic analysis. However, the relatively slow labeling kinetics of Caprola required high concentrations of fluorophore probe and relatively long labeling times, which limits its sensitivity and applicability, in particular *in vivo*. To address this limitation, we generated Caprola variants with up to 29-fold faster labeling rates than their predecessor. We demonstrate that our new Caprola variants record calcium transients in cells and in zebrafish larval brains under conditions where previous Caprola variants did not show labeling. We further expand the applicability of Caprola to activity-dependent marking of postsynaptic compartments, opening new avenues for coupling functional activity histories with downstream molecular and transcriptomic analyses.

## Introduction

Tracking neuronal activity with cellular resolution is essential for advancing our understanding of brain function. Powerful tools for tracking neuronal activities are calcium recorders, which translate a calcium transient into a permanent mark for later analysis^1^. These recorders separate the recording period from signal analysis, which makes them scalable with respect to the number of cells that can be analyzed and circumvents the technical challenges of imaging the brain of a freely moving animal^2^. One such approach exploits transcription of neuronal immediate early genes (IEGs), in which a calcium transient results in the expression of a reporter gene^3–6^. However, the relationship between neuronal activity and gene expression is only indirect^7^, and transcriptional recorders are limited in their temporal resolution and cell type compatibility. Optogenetic recording approaches like CaMPARI offer outstanding spatiotemporal resolution and enable direct recording of calcium transients, but their use is inherently constrained by the need for illumination^8–11^.

We recently introduced a strategy to record physiological activity based on a split HaloTag system^12, 13^. This <u>ca</u>lcium-dependent <u>prote</u>in <u>la</u>beling tool (Caprola) becomes irreversibly labeled in the presence of elevated calcium levels and a fluorescent probe, enabling *post hoc* analysis. Caprola enables direct recordings of successive periods of neuronal activity in cell culture and *in vivo*^12^. Further, the permanent nature of the fluorescence labeling allows sorting of heterogeneous cell populations for subsequent transcriptome analysis^12^. However, the early version of Caprola showed relatively slow labeling kinetics even in the presence of calcium, about 70-fold slower than the original HaloTag7^12, 14^. Its use in cells and neurons thus required concentrations of the fluorescent probe that are challenging to reach in animal models and relatively long recording times.

In this work, we present Caprola2 and Caprola3, optimized versions of Caprola. By incorporation of computationally designed elements and further rational engineering, we increased the labeling rate of Caprola2 and Caprola3 with fluorescent probes to approach the labeling rates of full-length HaloTag7. Caprola2 contains a fluorescent protein for ratiometric signal normalization, while Caprola3 is designed for applications where this fluorescent protein is not required. We demonstrate that the increased labeling rates of the new Caprola variants translate into a more sensitive recording of Ca^2+^ signaling in cells. Additionally, the increased labeling kinetics of Caprola2 enabled sensitive recordings of calcium transients at postsynaptic densities, extending calcium activity recording to the synaptic scale. Lastly, we demonstrate that Caprola2 outperforms Caprola1 in labeling neuronal activity in the zebrafish larval brain.

## Results

### Caprola engineering for faster calcium recording

Caprola is based on a split-HaloTag that renders the labeling activity of the protein with its chloroalkane (CA) fluorophore substrates dependent on the presence of Ca^2+ 12^. This split-HaloTag system comprises a truncated, circularly permuted HaloTag (cpHaloΔ), which maintains the overall fold of the protein but remains largely inactive, and a peptide (Hpep) that can reversibly bind to cpHaloΔ and restore its labeling activity. Caprola connects cpHaloΔ and Hpep through a linker comprising calmodulin (CaM) and M13 (Figure 1A). Complementation of split-HaloTag in Caprola occurs only when CaM binds M13 in a Ca^2+^-dependent manner.

**Figure 1.**
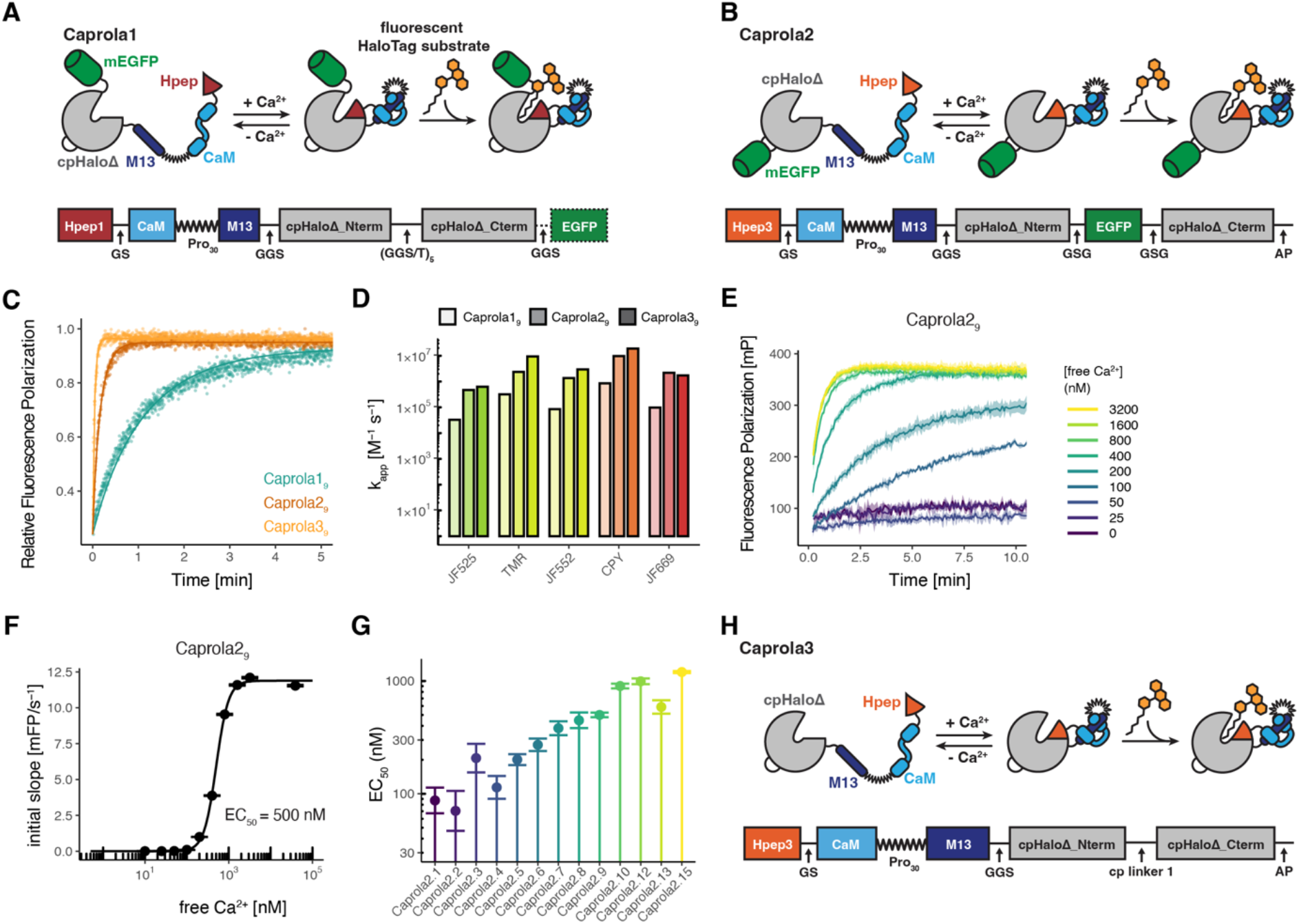
Design and characterization of the optimized Ca^2+^ recorders Caprola2 and Caprola3. (A) Scheme of Caprola1 design. (B) Scheme of Caprola2 design. (C) Labeling kinetics of Caprola1_9_, Caprola2_9_ or Caprola3_9_ (50 nM each) with TMR-CA (10 nM) in the presence of Ca^2+^ (5 mM) measured by fluorescence polarization (FP). (D) Apparent second-order rate constants (k_app_) of Caprola1_9_, Caprola2_9_ and Caprola3_9_ labeling with different fluorophore substrates in the presence of Ca^2+^. (E) Labeling kinetics of Caprola2_9_ (10 nM) with TMR-CA (2 nM) at different free Ca^2+^ concentrations measured by fluorescence polarization (FP). (F) Ca^2+^-sensitivity of Caprola2_9_ determined by plotting free Ca^2+^ concentrations against the initial slopes of the labeling reactions from (E). (G) Ca^2+^-sensitivity (EC_50_ values) of 13 Caprola2 variants determined as shown in (E) and (F). (H) Scheme of Caprola3 design.

To improve the sensitivity and temporal resolution of Caprola recordings, we sought to increase the labeling rate of the first-generation Caprola, in the following called Caprola1. The labeling kinetics of Caprola1 are almost two orders of magnitude slower than HaloTag7, which labels with apparent second order rate constants of k_app_ > 10^7^ M^−1^ s^−1^ for various rhodamine substrates^14^. HaloTag7 enables fast labeling in cells and has been used successfully *in vivo*^15–17^, thus faster Caprola labeling should facilitate applications in cells and *in vivo*.

First, we explored the impact of the termini of cpHaloΔ and Hpep in the first Caprola design. Guided by the effect of purification tags added to cpHaloΔ, we identified an extension at the C-terminal residue 141 of cpHaloΔ by the dipeptide AlaPro that increased the labeling rate by 1.8-fold (Figure S1A). Second, we investigated the effect of Hpeps with increased affinity for cpHaloΔ; these Hpeps were previously generated through computational approaches^12^. Different Hpeps were screened in the Caprola context for their effect on labeling kinetics in the presence and absence of Ca^2+^. We found that Caprola with Hpep3 (SKRDAREMFQAFRT, EC_50_ = 149 µM to cpHaloΔ1), which possesses a 20-fold higher affinity than Hpep1 (ARETFQAFRT, EC_50_ = 3.0 mM to cpHaloΔ1), had the highest labeling rate in the presence of Ca^2+^ without disproportionally increased background labeling in the absence of Ca^2+^ (Figure S1B). Last, to enable cytosolic Ca^2+^ activity recordings in cells, we added an N-terminal nuclear export signal (NES) sequence. Caprola1 has mEGFP fused to its C-terminus to normalize the fluorescence signal of the fluorescent probe to expression levels in individual cells (Figure 1A). However, a similar EGFP fusion to the improved Caprola variant introduced a Ca^2+^-dependent FRET (Figure S1C). Therefore, we placed mEGFP into the circular permutation linker of cpHaloΔ to both eliminate the unwanted FRET and conserve the faster labeling kinetics (Figure 1B, Figure S1C).

The resulting Caprola2 showed 7-fold faster labeling rates than Caprola1 (k_app_ = 2.4×10^6^ M^−1^ s^−1^ as compared to k_app_ = 3×10^5^ M^−1^ s^−1^) (Figure 1C). Compared to Caprola1 with mEGFP, Caprola2 melting temperatures in the presence and absence of Ca2+ were increased by 4.1 °C and 2.6 °C, respectively (Table S1). The labeling rate was increased for multiple spectrally distinct fluorescent probes (Figure 1D, Figure S2A), which have been previously used for the labeling of HaloTag7 in animal brains^13, 17^. The cooperative binding of four Ca^2+^ ions to CaM resulted in a sharp threshold of Ca^2+^ concentration below which Caprola2 labeling was minimal (Figure 1E, F). Furthermore, we introduced targeted mutations to M13 to yield Caprola2 variants Caprola2_1_-Caprola2_15_, exhibiting a spectrum of Ca^2+^ responsivities with EC_50_ values ranging from 70 nM to 1.2 µM (Figure 1G, Figure S3). All Caprola2 variants maintained high labeling rates upon Ca^2+^ binding and on average approx. 4,000-fold lower labeling rates in absence of Ca^2+^ (Figure S4, Table S2). For control experiments we also constructed constitutively active Caprola2_ON_, which comprised full-length cpHaloTag containing Hpep3, and constitutively inactive Caprola2_OFF_, which lacked the Hpep.

During the engineering work of Caprola, we found that removing the mEGFP further increased the labeling kinetics. To further improve both labeling rate and stability of cpHaloΔ in Caprola, we attempted to replace the mEGFP with a computationally designed linker connecting the original N- and C-termini of HaloTag7. We previously designed such linkers for cpHaloΔ using RosettaRemodel^18^. Following this approach, we generated and scored 1,500 structures by Rosetta, 500 each of linker length 21, 22 or 23 amino acids, to design a linker forming an α-helix to replace the flexible (GGS)^5^ linker of Caprola1^18^. The top 10 scoring linkers were implemented in cpHalo153/156, which were characterized for their thermostability and labeling kinetics. The most promising three linkers, further evaluated in the Caprola context, improved both thermostability and labeling rate (Figure S5, Table S1). One linker increased the melting temperature in the presence and absence of Ca^2+^ by 2.6 °C and 2.3 °C, respectively, and labeling kinetics in the presence of Ca^2+^ by 6-fold. Combining this linker with Hpep3 and the additional C-terminal dipeptide (AlaPro) of Caprola2, resulted in Caprola3 variant (Figure 1E), which exhibited a 29-fold increased labeling rate in the presence of Ca^2+^ compared to Caprola1 (k_app_ = 9.4×10^6^ M^−1^ s^−1^) and a 4-fold increased labeling rate compared to Caprola2 (Figure 1H), approaching the labeling rate of intact HaloTag7 with TMR-CA (k_app_ = 1.88×10^7^ M^−1^ s^−1 14^). In the absence of Ca^2+^, the labeling rate of Caprola3 was also increased (k_app_ = 925 M^−1^ s^−1^ for Caprola3 as compared to k_app_ = 38 M^−1^ s^−1^ for Caprola1, Figure S5). However, Caprola3 still possesses a ten-thousand-fold increase in labeling rate upon Ca^2+^ exposure.

### Caprola2 enables recording of Ca^2+^ transients at low probe concentrations

We then investigated the performance of Caprola2 in recording cytosolic changes in Ca^2+^ in immortalized mammalian U2OS cells. Caprola2 was used in these experiments for a ratiometric readout. We chose to compare variants with equivalent M13 peptides though most Caprola2 variants are less sensitive for Ca^2+^ than their Caprola1 counterpart (Table S2). U2OS cells expressing either Caprola1^6^ (EC_50_^Ca2+^ = 131 nM) or Caprola2_6_ (EC_50_^Ca2+^ = 271 nM) were incubated with low concentrations of a probe containing Janelia Fluor 669 (JF_669_-CA) (10 nM) in the presence or absence of thapsigargin, an inhibitor of the sarco(endo)plasmic reticulum Ca^2+^-ATPase (SERCA) pumps. Stimulation with thapsigargin led to strong labeling for Caprola2_6_, while only little signal was obtained for Caprola1_6_ (Figure 2A). In the presence of thapsigargin, fluorescence intensity ratios of Caprola2_6_ were 33-fold higher compared to Caprola1_6_ (Figure 2B). In the presence of EGTA, low labeling was observed for both Caprola versions (Figure 2A, B). In contrast to Caprola1_6_, Caprola2_6_ enabled to record changes in resting Ca^2+^ concentrations, as removal of extracellular Ca^2+^ with the Ca^2+^ chelator EGTA further reduced signal accumulation (Figure 2A, B). In flow cytometry assays, we evaluated the performance of Caprola2 variants with different calcium sensitivities, Caprola2_ON_ and Caprola2_OFF_ (Figure 2C). In these experiments, Caprola2_6_ showed the highest fold-change in median ratio (stimulated/basal) of 11.6 upon thapsigargin-treatment. For Caprola2_6_, fluorophore concentrations as low as 2 nM and labeling times as short as 2 minutes where sufficient to differentiate between treated and untreated cells (Figure S7). As expected, Caprola2_ON_ showed strong labeling independent of cellular treatment, while Caprola2_OFF_ showed negligible labeling under all conditions. A palette of spectrally distinguishable fluorescent probes enabled thapsigargin-induced labeling of Caprola2_6_ in cells, with JF_669_-CA, JF_646_-CA and SiR-CA showing the highest fold-range in median ratio (stimulated/basal) (Figure 2D, Figure S8).

**Figure 2.**
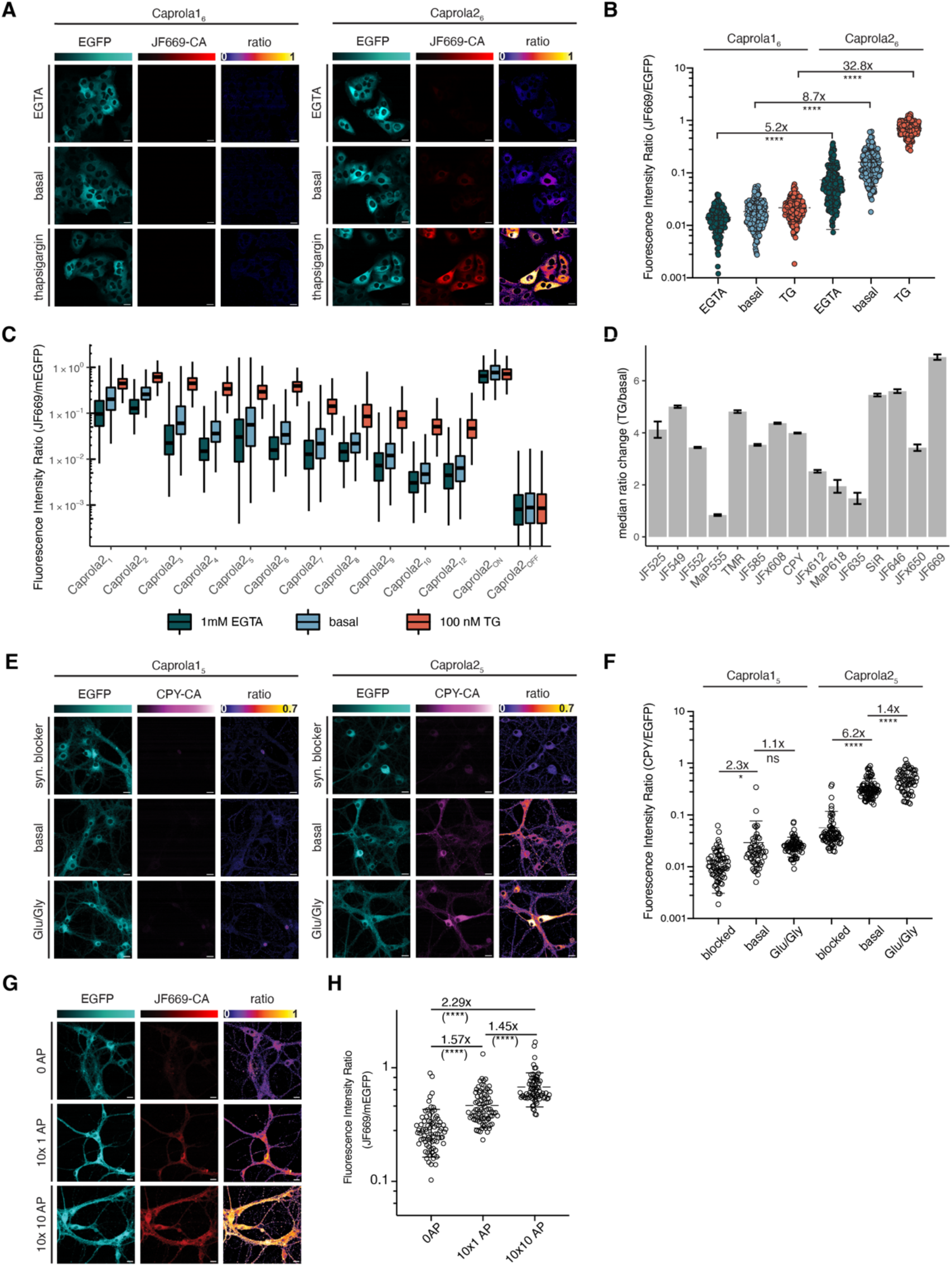
Caprola2 enables Ca^2+^ activity recordings at shorter time scale and with reduced fluorophore substrate concentrations. (A) Fluorescence images of U2OS cells expressing Caprola1_6_ or Caprola2_6_ incubated with JF_669_-CA (10 nM) in the presence or absence of thapsigargin (100 nM, 15 min). Ratio indicates JF_669_-CA/mEGFP. (B) Quantification of Caprola1_6_ and Caprola2_6_ labeling from experiment described in (A) (N > 170 cells, Welch’s t-test, ****: p ≤ 0.0001). (C) Flow cytometry analysis of U2OS cells expressing different Caprola2 variants incubated with JF_669_-CA (10 nM, 30 min) in the presence of thapsigargin (TG, 100 nM) or EGTA (1mM), or without treatment. Fluorescence intensity ratios (JF_669_-CA/mEGFP) are shown as box and whisker plots. (D) Flow cytometry analysis of U2OS cells expressing Caprola2_6_ incubated with different fluorophore substrates (10 nM, 15 min) in the presence or absence of thapsigargin (TG, 100 nM). Median fluorescence intensity ratio changes of stimulated/non-stimulated conditions are displayed. Error bars represent SEM. (E) Fluorescence images of primary rat hippocampal neurons expressing Caprola1_5_ or Caprola2_5_ labeled with CPY-CA (25 nM, 30 min) in the presence of synaptic blockers APV and NBQX (25 µM/10 µM) or glutamate and glycine (Glu/Gly; 10 µM/2.5 µM), or without treatment. Ratio indicates CPY-CA/mEGFP. (F) Quantification of Caprola1_5_ and Caprola2_5_ labeling from experiment described in (E) (N > 50 cells, Welch’s t-test, ns: p > 0.05, *: p ≤ 0.05, ****: p ≤ 0.0001). (G) Fluorescence images of primary rat hippocampal neurons expressing Caprola2_5_ incubated with JF_669_-CA (100 nM, 30 min pre-incubation with synaptic blockers, 10 min preincubation with fluorophore) upon defined electrical field stimulation. Action potentials (AP) were delivered in 10 short pulses (80 Hz) over the course of 6 min interspaced with resting periods (20 sec). (H) Quantification of Caprola2_5_ labeling from experiment described in (G) (N > 70 cells, Welch’s t-test, ****: p ≤ 0.0001). Panels (B, C, F, H): The center line indicates the median, and error bars indicate 25% and 75% quantiles. Panels (A, E, G): Scale bars: 20 µm.

Next, we investigated whether Caprola2 enabled recordings of cytosolic Ca^2+^ at nanomolar concentration of fluorescent probe in cultured neurons. We expressed Caprola1_5_ or Caprola2_5_ in primary rat hippocampal neurons, and labeled with only 10 nM CPY-CA (30 min) in the presence of different treatments. While silencing of synaptic transmission with APV and NBQX, antagonists of *N*-methyl-d-aspartate (NMDA) and α-amino-3-hydroxy-5-methyl-isoxazolepropionic acid (AMPA) receptors, respectively, resulted in decreased labeling relative to basally active neurons, stimulation with the neurotransmitters glutamate and glycine resulted in increased fluorescence intensity ratios (Figure 2E, F). These experiments demonstrate that Caprola2_5_ can record calcium transients evoked by different treatments at low nanomolar probe concentrations, while Caprola1_5_ labeling showed low fluorescence intensity over all conditions. In previous experiments, recording of neuronal activity with Caprola1_5_ required incubation with a 25-fold higher fluorophore concentration of 250 nM CPY-CA^12^. Caprola2_5_ proved more suitable than Caprola2_1_ or Caprola2_7_ in distinguishing the three different activity states (Figure S9). Compared to pharmacologically silenced neurons, Caprola2_5_ showed statistically significant differences in fluorescence labeling when increases in intracellular Ca^2+^ were elicited by only 10 electrically evoked, single action potentials, delivered over the course of 6 min in the presence of JF_669_-CA (100 nM) (Figure 2G, H).

### Improved Caprola enables recording calcium transients in excitatory postsynaptic densities

We applied Caprola2 to record Ca^2+^ activity histories in excitatory postsynaptic densities (Figure 3A). First, we exchanged mEGFP for mGreenLantern (mGL) due to its higher brightness in cells^19^ to compensate for the low expression levels of Caprola2 at postsynaptic densities. We chose to target Caprola2_5_ to the postsynaptic density protein 95 (PSD95) but avoided a direct fusion of Caprola2_5_ to PSD95, as overexpression of PSD95 has been reported to change function and morphology of dendritic spines^20^. Instead, we fused Caprola2_5_ to a genetically encoded intrabody against PSD95 (PSD95.FingR)^21^. To avoid a large fraction of unbound protein present in the cytosol, we employed a previously described targeted expression feedback loop to reduce cytosolic fluorescence (Figure 3B)^11, 21^. By fusing the zinc finger CCR5-L (ZF) and the transcriptional repressor KRAB(A) to the target protein, and incorporating a ZF binding site upstream of the promoter, the ZF-KRAB domain is presumed to recruit unbound protein to the nucleus. There, the ZF binds its target sequence, and KRAB(A) recruits the transcriptional repressor machinery to downregulate gene expression of the exogenous gene. As it has been discovered that the ZF binding site can be omitted^11^, we evaluated both expression systems in ectopically expressing HEK293T cells and cultured primary neurons. In HEK293T cells, we co-expressed the Caprola2_5_ fusion constructs either with a mitochondrially-localized PSD95-mScarlet construct or only mScarlet, and found co-localization at mitochondria only when PSD95 was expressed (Figure S10). In cultured neurons, we expressed the Caprola2_5_ fusion constructs with ZF binding site (Figure S11A, B) or without ZF binding site (Figure 3B, C) and found higher co-localization with PSD95 as well as higher fluorescence intensities for the variant lacking the ZF binding site (Figure S11C-E). Thus, we used this variant, in the following called postsyn-Caprola2_5_, for further experiments. We recorded fluorescence labeling of postsyn-Caprola2_5_ in cultured neurons (25 nM JF_669_-CA, 25 min) following different treatments: Stimulation with KCl increased fluorescence intensity ratios compared to basally active neurons, while silencing of neuronal transmission with APV/NBQX reduced fluorescence intensity ratios to levels as low as background signal in non-fluorophore treated neurons (Figure 3D, E). The fluorescence intensity ratios showed a pronounced heterogeneity in postsynaptic activity histories upon neuronal activation. By classifying postsynaptic densities into non-active (silent) and active subpopulations based on the interquartile range of the fluorescence intensity ratio, we found a median of ~71% of postsynaptic densities were active upon KCl treatment, as compared to only ~7 % under basal conditions, recorded over a time period of 25 minutes (Figure 3F). We further observed postsynaptic activity-dependent signal differences during long-term recordings of up to 90 min (Figure S12). In these experiments we used a reduced concentration of the fluorescent ligand (0.5 nM CPY-CA; Figure S12) to minimize the risk of progressive accumulation or saturation of labeling.

**Figure 3.**
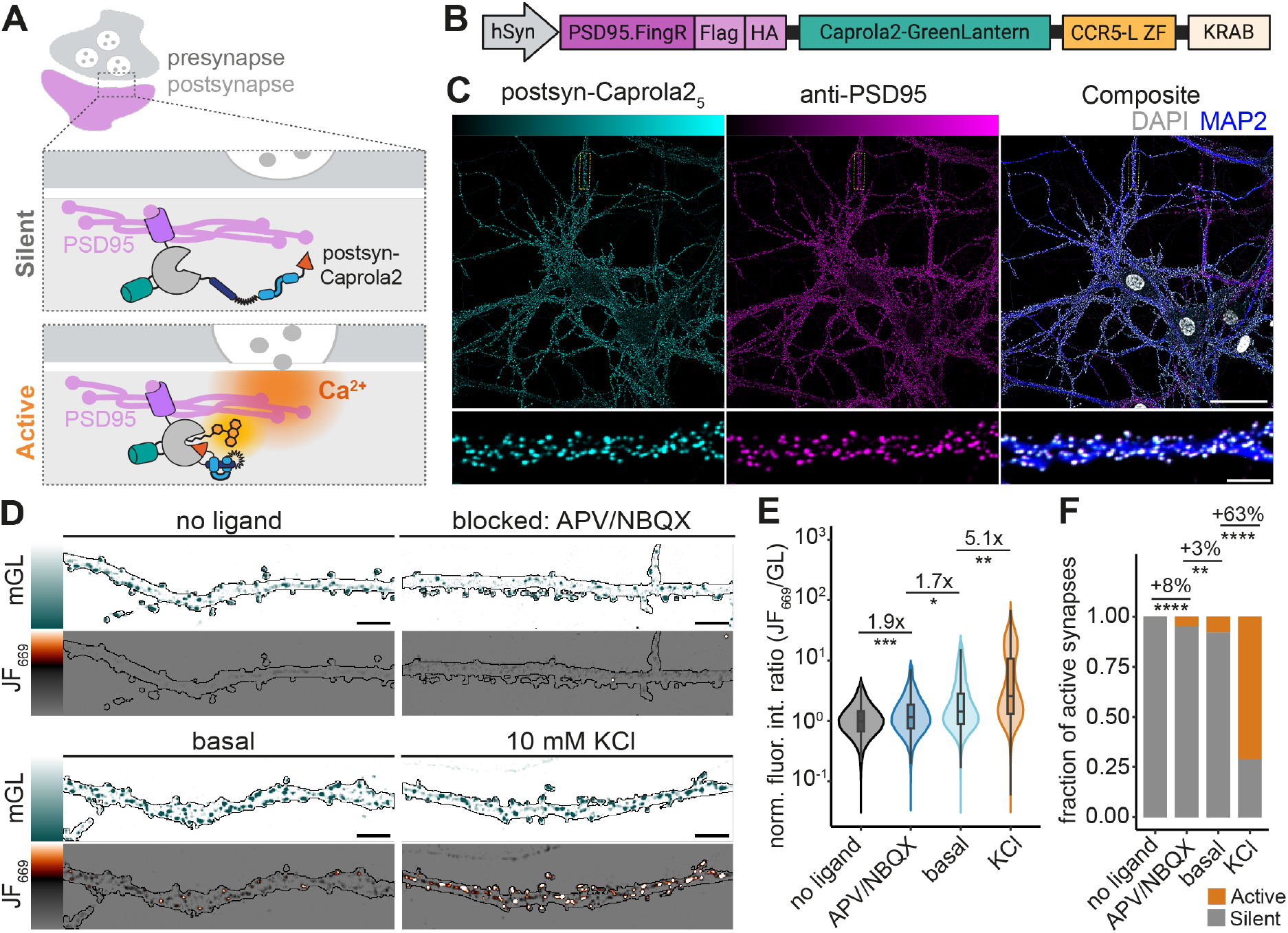
Caprola2 records Ca^2+^ activity in postsynaptic densities. (A) Concept of postsynaptic Ca^2+^ recordings with Caprola2. (B) Design of postsyn-Caprola2_5_ construct. Caprola2_5_ is targeted to postsynaptic densities via the PSD95 intrabody (PSD95.FingR). Expression is autoregulated using the CCR5-L zinc finger DNA-binding domain fused to the KRAB(A) transcriptional repressor to minimize excess unbound reporter. (C) Representative fluorescence images of primary rat hippocampal neurons expressing postsyn-Caprola2_5_. Left, GreenLantern fluorescence of postsyn-Caprola2_5_; middle, immunostaining for PSD95; right, merged image showing colocalization (white). MAP2 labels dendrites (blue) and DAPI labels nuclei (grey). Scale bars: 50 µm (overview) and 5 µm (magnified dendritic region). (D) Representative fluorescence images of dendritic regions following labeling with the HaloTag ligand JF_669_-CA (25 nM, 25 min) under the indicated conditions: synaptic blockade (APV/NBQX, 25 µM/10 µM), basal activity, or depolarization with KCl (10 mM). Non-fluorophore treated neurons indicate baseline signal. Images are displayed using identical acquisition and display settings. Scale bars, 5 µm. (E) Quantification of postsyn-Caprola2_5_ labelling from the experiments described in (D). Individual PSD95-positive synapses were quantified by calculating the JF_669_-CA/GreenLantern fluorescence ratio. Violin plots show the distribution of individual synapses, while boxplots indicate the median and interquartile range. Data were normalized to the median ratio of the no-fluorophore control. Statistical comparisons were performed using per-field-of-view summaries as the experimental unit (two-sided Wilcoxon rank-sum test with Benjamini– Hochberg correction). Data are from two independent experiments with two technical replicates each (91 fields of view). Ns, P > 0.05; *, P ≤ 0.05; **, P ≤ 0.01; ***, P ≤ 0.001. (F) Classification of postsynaptic sites according to their activity state under the treatment conditions described in (D). Thresholds for active versus inactive (silent) synapses were derived from JF_669_-CA /GreenLantern ratios in APV/NBQX-treated neurons, representing the inactive state, using an interquartile range (IQR)-based approach. Bar plots show the median fraction of synapses assigned to each activity class per field of view under the indicated experimental conditions.

### Caprola2 shows superior labeling performance *in vivo* compared to Caprola1

We previously used zebrafish lines pan-neuronally expressing Caprola1_1_ to record neuronal activation *in vivo* by simply incubating the fish with labeling substrates. However, substrate incubations for relatively long time periods were required to observe significant Caprola labeling. In freely swimming larvae incubated with 5 *μ*M CPY-CA, we observed labeling of Caprola1_1_ in the forebrain at the earliest after 30 min and in the hindbrain at the earliest after 60 min^12^. To investigate if the faster labeling kinetics of Caprola2 also translates into superior labeling performance *in vivo*, we generated zebrafish lines pan-neuronally expressing Caprola2_1_ and compared their labeling performance with the previously generated Caprola1_1_ zebrafish line^12^. Incubating freely-swimming zebrafish larvae expressing Caprola2_1_ with CPY-CA (5 µM) already resulted in very strong fluorescent labeling in the forebrain and significant labeling in the hindbrain after 30 min (Figure 4, Figure S13). The labeling intensity corresponded to the neuronal activity patterns observed during freely-swimming behavior^8^. In direct comparison, Caprola1_1_ labeling under the same conditions and microscope settings was minimal after 30 min and became detectable after 90 min only in the forebrain (Figure 4, Figure S14). The faster labeling kinetics of Caprola2 thus increases the temporal resolution of our recorder *in vivo*.

**Figure 4.**
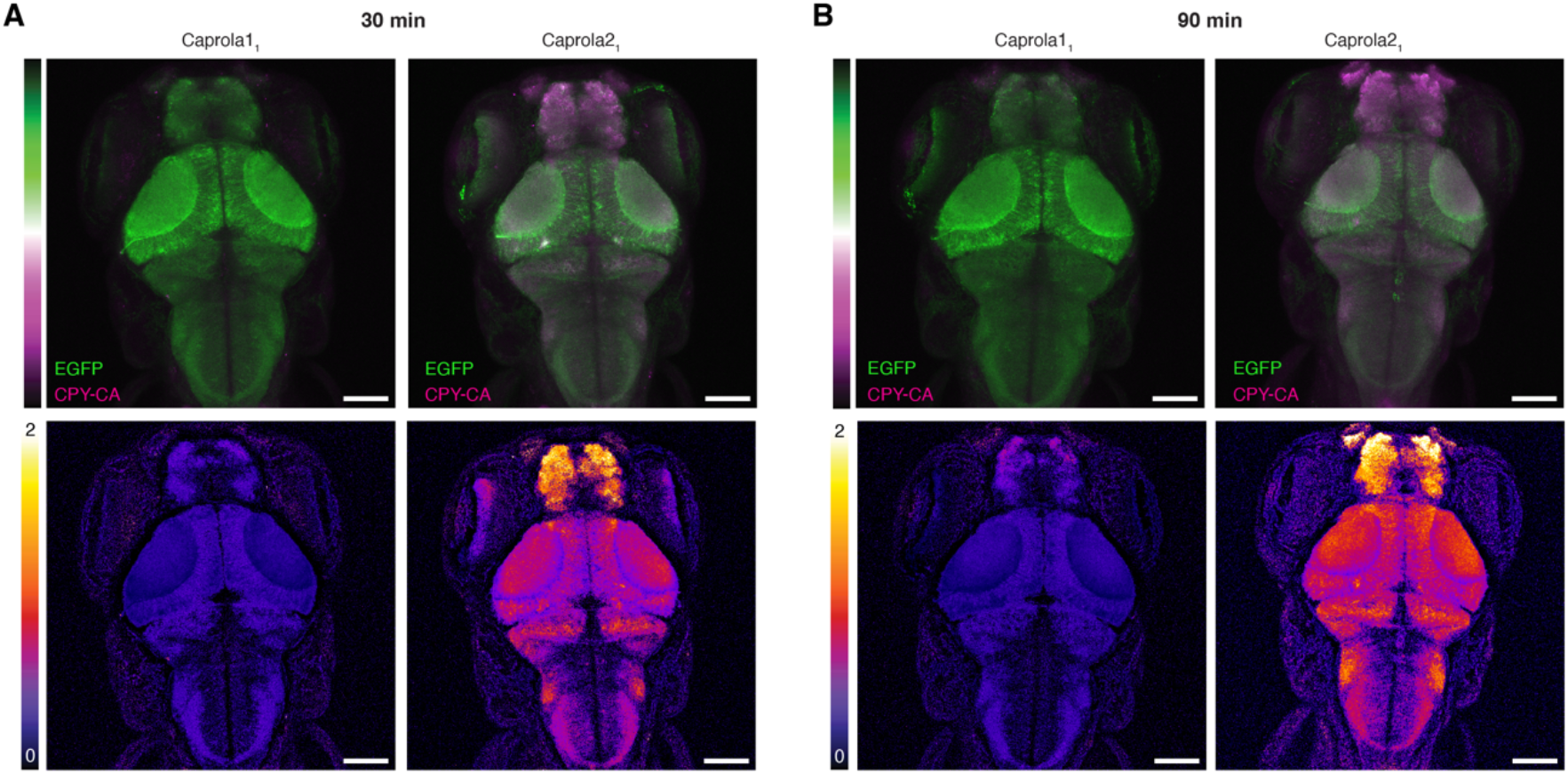
Caprola2 records neuronal activity in the zebrafish larval brain. (A) Representative fluorescence images of freely-swimming zebrafish larvae (4 dpf) pan-neuronally expressing Caprola1_1_ (left) or Caprola2_1_ (right) incubated with CPY-CA (5 µM) for 30 min (maximum intensity projections). The color scale bar represents the ratio CPY-CA/mEGFP of fluorescence intensities. (B) Representative fluorescence images of freely-swimming zebrafish larvae (4 dpf) expressing Caprola1_1_ (left) or Caprola2_1_ (right) incubated with CPY-CA (5 µM) for 90 min (maximum intensity projections). Ratio indicates CPY-CA/mEGFP. Scale bars: 100 µm.

## Discussion

We recently introduced the calcium-dependent protein labeling tool Caprola, which is based on the split-HaloTag system. Caprola converts transient Ca^2+^ elevations into permanent fluorescent marks for later analysis. The availability of spectrally distinguishable fluorescent substrates enables the sequential recording of activity periods within the same sample of individual animal. The effective temporal resolution of Caprola is determined by the speed of the labeling, the permeability of the fluorescent probe in cells and their pharmacokinetic properties in animals. While Caprola1 enables the recording of neuronal activity in zebrafish larvae and adult flies over time periods of tens of minutes, recordings at shorter timescales would require faster labeling or substrates with better pharmacokinetic properties. More efficient labeling would also allow to reduce the concentration of fluorescent probe, which reduces potential non-specific background signal.

In this work, we focused on increasing the performance of Caprola by increasing its labeling speed. We developed Caprola2 and Caprola3, which react 7-fold and 29-fold faster than Caprola1 with fluorescent HaloTag substrates, respectively. For both Caprola2 and Caprola3, the labeling reaction is increased in the presence and absence of Ca^2+^, such that the overall dynamic ranges of Caprola2 and Caprola3 did not change compared to Caprola1. Importantly, the substantially faster labeling rates directly translate into more efficient labeling in cells. In comparison to Caprola1, the faster labeling rates of Caprola2 enabled reduction of recording intervals in cells to few minutes and fluorophore substrate concentrations down to 2 nM. In neurons, this increased sensitivity of Caprola2 allowed the detection of calcium activity corresponding to as few as approximately ten action potentials, substantially extending the range of detectable physiological signals. Caprola2 incorporates an mEGFP that enables ratiometric normalization of labeling signals, and a set of Caprola2 variants with different Ca^2+^ sensitivities provides flexibility for adjusting to diverse experimental requirements. Caprola3 does not contain mEGFP for normalization, but exhibits even faster labeling kinetics than Caprola2, approaching the labeling speed of HaloTag7. Therefore, Caprola3 should become useful for applications that either do not require signal normalization or can achieve normalization through other means. Given that Caprola3 almost approaches the labeling speed of HaloTag7, further improvements in cellular labeling speed might require the design of probes that are more cell-permeable and present more favorable pharmacokinetic properties rather than additional protein engineering.

To demonstrate the utility of our new Caprola variants, we targeted Caprola2 to the postsynaptic scaffold protein PSD95 at its endogenous expression levels to record Ca^2+^ activity histories in postsynaptic densities. Under our experimental conditions, we detected specific post-synaptic fluorescent labeling of individual synapses upon stimulation, establishing that the increased sensitivity of Caprola2 allows recordings of spatially confined calcium transients at synapses. Beyond synaptic labeling, the permanent nature of the Caprola2 signal offers the potential to link local calcium activity histories to downstream molecular readouts, including transcriptomic analyses to investigate molecular mechanisms underlying synaptic heterogeneity. The recording of calcium transients in neuronal synapses illustrate Caprola’s broader potential for applications within defined subcellular compartments. *In vivo*, Caprola2 showed substantially higher labeling activity compared to Caprola1, allowing the recording of neuronal activity in freely-swimming zebrafish larvae in both fore- and hindbrain in about 30 min. This increased sensitivity of Caprola2 *in vivo* should facilitate the recording of neuronal activation for *post hoc* analysis under a wide range of conditions.

In summary, we have introduced variants of Caprola, Caprola2 and Caprola3, with substantially improved labeling rates relative to their predecessor. The faster labeling enables calcium recordings over shorter time periods and at lower substrate concentrations, both in cell culture and *in vivo*. We anticipate Caprola2 and Caprola3 to become important tools to record physiologically relevant calcium transients for their correlation with biological phenotypes.

## Acknowledgments and funding

This work was supported by the Max Planck Society and École Polytechnique Fédérale de Lausanne (EPFL). The authors thank B. Koch, A. Bergner, J. Kress, P. Breuer, A. Herold, B. Réssy, and D. Schmidt (all MPIMR) for providing reagents or material, and J. Keith for preliminary experiments. We thank M. Tarnawski (protein expression and characterization facility, MPIMR), S. Fabritz, T. Rudi, and J. Kling (all mass spectrometry facility, MPIMR) for their support. We thank Yuqi Zhang for critical reading of the manuscript. N. Porzberg was supported by the European Union’s Horizon 2020 grant 955623, J. Wilhelm by the Max Planck School Matter to Life, J. Heck by a fellowship from the EMBL Interdisciplinary Postdoc (EIPOD) program under Marie Sklodowska-Curie Actions, the EIPOD4 COFUND IV Grant (grant agreement number 847543) as well as the European Molecular Biology Laboratory. N. Porzberg was supported by the Heidelberg Biosciences International Graduate School (HBIGS).

