## Supporting Information for "Fast calcium-dependent fluorescent labeling for recording of neuronal activation"

### Materials and Methods

#### Chemicals

Chemicals and reagents were purchased from the following commercial providers: Merck KGaA, Honeywell International Inc., Fisher Scientific International Inc., Carl Roth GmbH & Co. KG, VWR International, Acros Organics B.V.B.A.

Fluorescent substrates for HaloTag were synthesized according to published procedures[1-4], purchased from Promega Corporation or were kind gifts from Dr. L. D. Lavis (Janelia Research Campus). Their chemical structure and photophysical properties are reported in Figure S15.

#### Biochemistry and molecular biology

General information

DNA and protein concentrations were determined with a NanoDrop 2000c spectrometer (Thermo Fisher Scientific) by measuring absorption at 260 nm and 280 nm, respectively. DNA solutions were stored at -20 °C and proteins were mixed 1:1 with 90% (w/v) glycerol in activity buffer (Method Table 1) and stored at -20 ºC. Microplate reader experiments were performed using a microplate reader Spark20M instrument (Tecan Group). Data analyses of *in vitro* experiments were performed using custom R scripts[5,6]. All buffer compositions are summarized in Method Table 1.

Design of circular permutation linkers

A cpHaloTag structural model was generated from the HaloTag crystal structure (PDB: pdb_00004kaf, chain A). Residues 1–13, 306, and 307 were removed, and residue 153 was deleted to expose new termini. The sequence was then renumbered to match the cpHaloTag construct. The resulting model was relaxed with Rosetta FastRelax[7] using the following command:

```
$ROSETTA3/bin/relax.mpi.linuxgccrelease  \
    -in:file:s input/cpHalo.pdb \
    -nstruct 240 \
    -relax:constrain_relax_to_start_coords \
    -relax:ramp_constraints false \
    -ex1 \
    -ex2 \
    -use_input_sc \
    -flip_HNQ \
    -no_optH false \
    --out:path:all output_relax
```

The lowest-scoring relaxed model served as the template for cp-linker design with RosettaRemodel[8]. Three blueprint files were prepared, keeping the backbone fixed except for positions flanking the cp-linker, and introducing helical linkers of 21-23 amino acids. An excerpt of a blueprint file is shown below:

```
[…]
149 T .
150 L .
151 E .
152 I L PIKAA I
0 X L ALLAAxc
0 X L ALLAAxc
0 X L ALLAAxc
0 X H ALLAAxc
0 X H ALLAAxc
[…]
0 X H ALLAAxc
0 X H ALLAAxc
0 X L ALLAAxc
0 X L ALLAAxc
0 X L ALLAAxc
153 I L PIKAA I
154 G .
155 T .
156 G .
[…]
```

RosettaRemodel was run with the following command:

```
$ROSETTA3/bin/remodel.mpi.linuxgccrelease
    -in:file:s input/cpHalo_relaxed.pdb
    -remodel:blueprint input/cpHalo.remodel
    -run:chain A
    -remodel:num_trajectory 10
    -nstruct 10000
    -save_top 100
    -ex1
    -ex2
    -out:path:all output_remodel
    -out:file:scorefile score.sc
    -no_optH false
    -find_neighbors
```

```
    -design_neighbors
```

For each linker length, top-scoring designs were selected for experimental evaluation in the cpHalo153/156 context. This included three linkers of 21-residue length, four linkers of 22-residue length and three linkers of 23-residue length. For the 21-residue linkers, three designs were chosen, however, the second top-ranked model was excluded due to a slight kink in the helix. For the 23-residue linkers, three designs were chosen, however, the top-ranked model was excluded due to a pronounced kink in the helix, and the 4th-ranked model was replaced with the 5th-ranked design to sample more diverse helix placements. The chosen designs were evaluated in the cpHalo153/156 context for thermostability and labeling kinetics. The 3 top performing variants in the cpHalo153/156 context, one 21-residue linker and two 23-residue linkers, were further evaluated in the Caprola context.

Molecular cloning

Molecular cloning was done using either Gibson assembly[9] or the Q5 site-directed mutagenesis kit (NEB) according to manufacturer's protocol. Polymerase chain reaction amplifications were performed using the KOD Hot Start DNA Polymerase master mix (Sigma-Aldrich) according to manufacturer's protocol. Plasmids generated by Gibson assembly were chemically transformed in *E. coli* strains E.cloni 10G (Lucigen) or NEB stable (NEB) for pAAV-hSyn1 vectors and pTol2-*elavl3*(HuC) vectors. Sequences were verified by Sanger sequencing (Eurofins Scientific or Microsynth Seqlab) with particular attention to the ITR integrity on pAAV-hSyn1 vectors and the Tol2 recombination sites on pTol2-*elavl3* vectors.

Recombinant protein expression and purification from *E.Coli*

Proteins were expressed in *E. coli* strain BL21(DE3)-pLysS (Novagen). Lysogenic broth (LB) cultures were grown at 37 °C until reaching an optical density at 600 nm ($OD_{600}$) of 0.8. Transgene expression was induced by the addition of 0.5 mM isopropyl-β-D-thiogalactopyranoside (IPTG) and cultures were grown at 16 °C overnight. Cells were harvested by centrifugation (15 min, 4 500 g, 4 °C), resuspended in IMAC lysis buffer (Method Table 1) and lysed by sonication on ice (5 min, 5 cycles, 70%). Lysates were cleared by centrifugation (20 min, 75 000 g, 4 °C) and proteins were purified via immobilized metal affinity chromatography (IMAC) using a HisTrap FF crude column (Cytiva) on an ÄktaPure FPLC system (Cytiva). Buffer exchange to activity buffer (Method Table 1) was conducted using a HiPrep 26/10 desalting column (Cytiva).

Proteins were concentrated using Amicon® Ultra-15 centrifugal filter devices (Merck) with a molecular weight cut-off (MWCO) smaller than the protein size to a final concentration of 100-500 µM. Correct size and purity of proteins were assessed by SDS-PAGE and high-resolution mass spectrometry (HRMS). Protein sequences are listed at the end of the supplementary information.

Protein thermostability measurement via nanoDSF

Thermostabilities of proteins were measured with 16 µM protein solution in activity buffer (Method Table 1) on a Prometheus NT 48 nanoscale differential scanning fluorimeter (NanoTemper) over a temperature range from 20 °C to 95 °C with a heating rate of 1 °C $min^{-1}$. Changes in the ratio of the fluorescence intensities at 350 nm and 330 nm were monitored. Indicated melting temperatures (mean of technical duplicates) correspond to the point of inflection (maximum first derivative).

Labeling kinetics of Caprola2 and Caprola3

Labeling kinetics of Caprola2 and Caprola3 variants were measured by recording fluorescence polarization over time in a microplate reader at 37 °C using black, non-binding, flat bottom, 96-well, polystyrene plates (OptiPlate, PerkinElmer) with a final reaction volume of 200 µL. Measurements were performed in FP buffer (Method Table 1) in technical triplicates. Final concentrations of 50 nM protein and 10 nM TMR-CA were used. Protein and substrate were prepared in 100 µL and the reactions were started by adding 100 µL 10 mM $CaCl_2$ in FP buffer (Method Table 1) (5 mM final concentration). Control experiments were conducted where 100 µL of buffer without $CaCl_2$ was added. A second-order reaction rate equation (equation 1) was fit to the data to obtain estimates for the apparent second-order rate constant $k_{app}$. $FP_{free}$ was fixed to the FP of the free dye in buffer. Uncertainties and confidence intervals of fitted parameters were estimated using the Monte Carlo method[10] (N = 1000).

$$(1) \qquad FP(t) = FP_{bound} + \frac{FP_{free}-FP_{bound}}{[A]_0} \cdot \frac{[A]_0([A]_0-[B]_0)e^{([A]_0-[B]_0)k_{app}\cdot t}}{[A]_0 e^{([A]_0-[B]_0)k_{app}\cdot t}-[B]_0}$$

with:

| | |
|---|---|
| t: | time |
| FP(t): | FP at time t |
| $FP_{bound}$: | FP of the bound dye |
| $FP_{free}$: | FP of the free dye |
| $[A]_0$: | dye concentration at t = 0 |
| $[B]_0$: | protein concentration at t = 0 |
| $k_{app}$: | apparent second-order rate constant |

To measure the reaction speed of the background labeling reaction in absence of $CaCl_2$, which is orders of magnitudes slower, fluorescence polarization assays were conducted as described above without adding $CaCl_2$. To measure over 24 h, a humidity cassette and a much lower sampling rate (1/900s) were used to limit. Data analysis was performed as described above, but in cases where FP did not plateau after 24 h, $FP_{bound}$ was fixed to the average plateau reached after full labeling of Caprola in presence of $Ca^{2+}$.

Labeling kinetics of $Caprola2_9$ and $Caprola3_9$ with different HaloTag substrates

Labeling kinetics of $Caprola2_9$ and $Caprola3_9$ with various HaloTag substrates were measured by recording fluorescence polarization over time in a microplate reader at 37 °C using black, non-binding, flat bottom, 96-well, polystyrene plates (OptiPlate, PerkinElmer) with a final reaction volume of 200 µL. Measurements were performed in FP buffer (Method Table 1) supplemented with 200 µM EGTA in technical triplicates. Final concentrations were 50 nM protein and 10 nM HaloTag substrate (TMR-CA, CPY-CA, $JF_{669}$-CA, $JF_{552}$-CA or $JF_{525}$-CA). Protein and substrate were prepared in 100 µL and the reactions were started by adding 100 µL 10 mM $CaCl_2$ in FP buffer (Method Table 1) (5 mM final concentration) using the injector module of the microplate reader. A second-order reaction rate equation (equation 1) was fit to the data to obtain estimates for the apparent second-order rate constant $k_{app}$. $FP_{free}$ was fixed to the FP of the free dye in buffer. Uncertainties and confidence intervals of fitted parameters were estimated using the Monte Carlo method[10] (N = 1000).

$Ca^{2+}$-dependence of Caprola2 labeling

The sensitivity of Caprola2 variants for calcium ($EC_{50}$) was determined by measuring labeling kinetics at different free $Ca^{2+}$ concentrations. Kinetics were recorded by measuring fluorescence polarization over time in a microplate reader at 37 °C using black, non-binding, flat bottom, 96-well, polystyrene plates (OptiPlate, PerkinElmer) with a final reaction volume of 200 µL. A calcium calibration kit (Thermo Fisher Scientific) was used to obtain buffers with precise and buffered free $Ca^{2+}$ concentrations ranging from 0 µM to 39 µM (100 mM KCl, 30 mM MOPS, pH 7.2 and $K_2$EGTA / CaEGTA in different ratios). Reactions were initiated by mixing equal volumes of Caprola2 protein (10 nM final) and TMR-CA (2 nM final) substrate in the respective buffers. Data from triplicates were averaged and a second-order reaction rate equation (equation 1) was fit to the data when at least 200 mFP were reached to obtain estimates for the apparent second-order rate constant $k_{app}$. The initial slope at time = 0 ($s_{t=0}$) was calculated using the derivative of equation 1 at t = 0 (equation 2).

(2) $$s_{t=0} = \frac{dFP}{dt}(t=0) = [B]_0 \cdot k_{app}(FP_{free} - FP_{bound})$$

with:

| | |
|---|---|
| t: | time |
| $s_{t=0}$: | initial slope at t = 0 |
| $FP_{bound}$: | FP of the bound dye |
| $FP_{free}$: | FP of the free dye |
| $[B]_0$: | protein concentration at t = 0 |
| $k_{app}$: | apparent second-order rate constant |

In conditions where 200 mFP were not reached due to the reaction being too slow, a linear model (equation 3) was fit to the data to determine the initial slope $s_{t=0}$.

(3) $$FP(t) = FP_{free} + t \cdot s_{t=0}$$

with:

| | |
|---|---|
| t: | time |
| $s_{t=0}$: | initial slope at t = 0 |
| $FP_{free}$: | FP of the free dye |

Initial slopes ($s_{t=0}$) were plotted against the free $Ca^{2+}$ concentration and a sigmoidal dose response model (equation 4) was fit to this data to estimate $EC_{50}$ values. Uncertainties and confidence intervals of fitted parameters were estimated using the Monte Carlo method[10] (N = 1000).

(4) $$s_{t=0}([Ca^{2+}_{free}]) = \frac{s_{max}}{1+10^{(LogEC50 - log10([Ca^{2+}_{free}]))\cdot HillSlope}}$$

with:

| | |
|---|---|
| $s_{t=0}$: | initial slope at t = 0 |
| $s_{max}$: | maximal initial slope at t = 0 |
| $[Ca^{2+}_{free}]$: | free $Ca^{2+}$ concentration |
| LogEC50: | $log_{10}$ of the half maximal effective concentration ($EC_{50}$) |
| HillSlope: | Hill coefficient |

### Cell biology

General information

Mammalian cells were cultured in high-glucose (4.5 g $L^{-1}$) and pyruvate (110 mg $L^{-1}$) containing Dulbecco's Modified Eagle Medium (DMEM)+GlutaMax™ (Gibco) medium supplemented with 10% fetal bovine serum (FBS) and phenol-red (herein termed DMEM) at 37 °C in a humidified incubator with 5% $CO_2$ atmosphere, and routinely passaged every 2–3 days. Primary rat hippocampal neurons were cultured in NeuroBasal medium supplemented with 1x GlutaMax™, 1x B-27™, and 1x Pen/Strep™ (all Gibco) at 37 °C in a humidified incubator with 5% $CO_2$ atmosphere. Primary mouse hippocampal neurons were cultured in

NeuroBasal medium supplemented with 5 mM L-glutamine and 2% B-27™ (both Gibco) at 37°C in a humidified incubator with 5% $CO_2$ atmosphere.

Generation of stable cell lines

Stable cell lines were prepared using the Flp-In™ T-REx™ System (Thermo Scientific). U2OS cells were transfected at 60% confluency using Lipofectamine™ 3000 (Thermo Scientific) according to manufacturer's protocol using a mix of 440 ng pDNA5/FRT/TO plasmid of interest mixed with 3560 ng pOG44 plasmid encoding the FlpIn recombinase. After 6 h incubation, the medium was replaced and cells were grown for additional 24 h. Cells were selected for 72 h at 37 °C in a humidified 5% $CO_2$ incubator using DMEM supplemented with 100 µg $mL^{-1}$ Hygromycin B (Gibco). Surviving cells were recovered in DMEM without Hygromycin B until cell confluency of 80% was reached. Highly expressing cells according to mEGFP fluorescence intensity were sorted using a FACS Melody (Becton Dickinson, excitation 488 nm, filter 530/30 nm). Stably expressing cells were maintained as explained above or stored in freezing medium (DMEM + 5% DMSO) at -80 °C or in the liquid nitrogen tank (vapor phase). Cell lines were regularly tested for mycoplasma contamination and were mycoplasma-free.

Transient transfection of cells

HEK293T cells were seeded onto 12-well plates and grown to 30–40% confluency. Cells were co-transfected with pairs of plasmids of interest in a 1:1 ratio using FuGENE® 6 (Promega) according to manufacturer's protocol. After 6 h incubation, medium was replaced and cells were grown for additional 24 h.

Chemical fixation of cells

After experimental treatment, cells were washed twice with pre-warmed 1x PBS (Gibco) prior to fixation. Freshly prepared and pre-warmed (37 °C) 4% Paraformaldehyde dissolved in 1x PBS without $Mg^{2+}$/$Ca^{2+}$ was applied for 15-20 min at room temperature. Cells were washed three times for 15 min with 1x PBS (Gibco) and kept at 4 °C for up to one week.

Recording intracellular calcium levels in mammalian cells with Caprola2

*Chemical activation of U2OS cells.* Stably expressing U2OS cells were seeded onto 96-well culture dishes and grown to 80% confluency. $Ca^{2+}$ modulators (100 nM thapsigargin or 1 mM EGTA) and fluorophore substrates in indicated concentration in DMEM were simultaneously applied and cells were incubated at 37 °C in a humidified incubator with 5% $CO_2$ atmosphere for the indicated time interval (0.5 min – 30 min). Cells were then washed with DMEM supplemented with HaloTag7 protein (1 µM) for 5 min, followed by two washes with pre-warmed 1x PBS (Gibco). Cells were subsequently analyzed by either flow cytometry or fluorescence microcopy.

Flow cytometry

After treatment, U2OS cells were detached from dishes using transparent TrypLE™-Express (Gibco, 1/4 of dish volume) at 37 °C in a humidified 5% $CO_2$ incubator for 5 min. Cells were re-suspended in 1x PBS containing 2% FBS and transferred into a flow cytometry compatible dish (non-adhering 96-U-well, FALCON). Samples were subjected to the autosampler of a Fortessa X-20 flow cytometer (BD Biosciences). Cell populations were gated for live (SSC-A/FSC-A) and single cells (FSC-H/FSC-A). Fluorophores were recorded as follows: mEGFP (488 nm excitation, 530/30 nm emission), fluorophores in the range from 525 nm to 570 nm (561 nm excitation, 575/26 nm emission), fluorophores in the range from 580 nm to 620 nm (561 nm excitation, 610/20 nm emission) and fluorophores in the range from 620 nm to 680 nm (648 nm excitation, 660/20 nm emission). Photomultiplier tube detectors were adjusted to avoid signal saturation.

*Flow cytometry data analysis.* Raw data from quantitative flow cytometry measurements were imported into the FlowJo suite (BD). First, live (SSC-A/FSC-A) and single cell (FSC-H/FSC-A) gates were adjusted and cells displaying fluorescence intensities of mEGFP below $10^3$ a.u. were excluded from analysis. Fluorescence intensity ratios were calculated for every cell individually by dividing the fluorescence intensities from fluorophores by fluorescence intensities from mEGFP. Dot plots showing whole populations or ratio density plots were created in FlowJo's layout manager. Quantitative assessment and statistical analysis were performed using custom R scripts[5].

Confocal microscopy

Confocal microscopy of U2OS cells and primary cultured rat hippocampal neurons cytosolically expressing Caprola was either performed on a commercial Leica SP8 or Stellaris 5 inverted microscope, both equipped with a white line laser (WLL) and hybrid photodetectors for single molecule detection (HyD SMD detector). Live cell imaging was performed at 37 °C with a 5% $CO_2$ atmosphere in a humidified chamber. Fixed samples were allowed to equilibrate to 37 °C for 15 min to avoid thermal drifting during image acquisition. The following settings were used for image acquisition: 20x/0.80 air objective; image size: 581.82 x 581.82 µm; scan speed 600-700 MHz; pinhole 1-2 airy units; line averages (4-8) and 8-bit or 16-bit depth. Dual-color imaging was done sequentially to prevent bleed-through.

Imaging data analysis

Image analysis was performed with FIJI[11]. ROIs were hand-segmented and mean fluorescence intensities from individual ROIs were derived for multiple fields-of-views. Cells displaying unhealthy phenotypes were excluded from analyses. Brightness and contrast were adjusted if needed, and scale bars were added for image representation. Ratiometric images were generated by using the BRET analyzer plug-in[12]. In short, image stacks were split into their individual fluorescence channels. The mEGFP channel was used for thresholding (Chastagnier method, default settings) and divisor of ratiometric fluorescence intensities (fluorescent substrate/mEGFP). A Gaussian blur was applied to resulting ratiometric images, which are represented with the 'fire' look-up table.

Postsyn-Caprola2 localization in HEK293T cells

*Immunofluorescence staining.* HEK293T cells transiently expressing the constructs of interest were washed with 1x PBS (Gibco), fixed with 4% pre-warmed PFA (20 min) and washed thrice with 1x PBS (Gibco) for 15 min. Then, cells were permeabilized with 0.3% Triton X-100/1x PBS for 3 min at ambient temperature, and washed thrice with washing buffer (Method Table 1) for 10 min. Samples were incubated with primary antibodies diluted in washing buffer for 1h at ambient temperature: camelid sdAB FluoTag-X4 anti-GFP labeled with ATTO488 (NanoTag Biotechnologies, diluted 1:300) and camelid sdAB FluoTag-X4 anti-RFP labeled with AZDye568 (NanoTag Biotechnologies, diluted 1:300). DAPI (Thermo Scientific) was added to the antibody incubation step at a final concentration of 0.1 $\mu$g $mL^{-1}$. After three washing steps with 1x PBS (Gibco) for 10 min, cells were stored in 1x PBS at 4 °C and imaged within one week after staining.

*Image acquisition and image analysis.* Spinning disk confocal microscopy of HEK293T cells was performed on a Nikon Ti2-E inverted microscope coupled to a Crest X-Light V2/V3 spinning disk confocal module (Nikon). The microscope was further equipped with a CFI Plan Apo Lambda D 60X Oil/1,42/0,15 objective (Nikon), Celesta light engine (Lumencor), and a Kinetix 10.2 MP Mono Back Illuminated sCMOS camera (Photometrics). Images were acquired as Z-stacks of up to 70 planes at 0.2 $\mu$m steps size. Maximum projections were generated for the multi-channel images, background was subtracted using the rolling ball algorithm, and brightness and contrast levels were scaled equally for images across conditions using FIJI[11].

Recording somatic and post-synaptic calcium levels in primary neurons with Caprola2

*Recombinant adeno-associated virus (rAAV) preparation*. rAAVs were prepared as previously described[13]. In brief, plasmids pRV1 (AAV2 Rep and Cap sequences), pH21 (AAV1 Rep and Cap sequences), pFD6 (adenovirus helper plasmid) and the AAV plasmid containing the recombinant expression cassette driven by hSyn1 promoter and flanked by AAV2 inverted terminal repeats (ITRs) were transfected via Polyethylenimine 25000 (Sigma Aldrich) or Lipofectamine™ 3000 (Thermo Scientific) into HEK293 cells. After 3-5 days of post transfection, cells were harvested and lysed using TNT extraction buffer (20 mM Tris pH 7.5, 150 mM NaCl, 1% TX-100, 10 mM $MgCl_2$). Cell debris was removed by centrifugation and supernatants were treated with Benzonase. rAAVs were purified from excess medium and cell supernatant via FPLC using AVB Sepharose columns and subsequently concentrated using centrifugal filter devices (Amicon, Merck KgaA) with a MWCO of 100 kDa, followed by buffer exchange to PBS pH 7.3. Titers of rAAVs were quantified by qPCR as previously described[14].

*Preparation of primary rat hippocampal neuron cultures and rAAV transduction*. Neurons were obtained from isolated hippocampi from newborn rats (WISTAR) as described previously[15]. Procedures were performed in accordance with the Animal Welfare Act of the Federal Republic of Germany (Tierschutzgesetz der Bundesrepublik Deutschland, TierSchG) and the Animal Welfare Laboratory Animal Regulations (Tierschutzversuchsverordnung). According to the TierSchG and the Tierschutzversuchsverordnung, no ethical approval from the ethics committee is required for the procedure of euthanizing rodents for subsequent extraction of tissues. The procedure for euthanizing rats was supervised by animal welfare officers of the Max Planck Institute for Medical Research (MPIMR) and conducted and documented according to the guidelines of the TierSchG (permit number assigned by the MPIMR: MPI/T-35/18). Neurons were dissociated by tryptic digestion and seeded in 24-well or 96-well glass bottom plates coated with poly-L-ornithine (100 µg $mL^{-1}$ in $ddH_2O$) and laminin (1 µg $mL^{-1}$ in 1x HBSS). Neurons were cultured for 7 days before rAAV-mediated transduction. Equivalents of $10^9$ –$10^{10}$ GC $mL^{-1}$ of purified rAAVs (serotype 2/1) were applied to individual samples and cultures were allowed to express transgenes for 7-11 days.

*Preparation of primary mouse hippocampal neuron cultures and rAAV transduction*. Mouse hippocampal neurons were obtained from isolated hippocampi from newborn mouse pups P0-P1 (C57BL/6). Procedures were performed in accordance with the Animal Welfare Act of the Federal Republic of Germany (Tierschutzgesetz der Bundesrepublik Deutschland, TierSchG) and Animals were sacrificed according to the local government (Rheinland-Palatinate, Germany) regulations for animal welfare. The use of animals was approved based on a grant under the reference number 41a/177-5865-§11 ZVTE 30.04.2014, by the district administration of Mainz-Bingen. Hippocampal neurons were dissociated by trypsin-EGTA for 15 min and seeded at a density of 70 000 cells per ø18 glass coverslips in 12-well plates coated with poly-L-lysine. Neurons were cultured for 7 days before rAAV-mediated transduction. Equivalents of $10^9$–$10^{10}$ GC $mL^{-1}$ of purified rAAVs (serotype 2/1) were applied to individual samples and cultures were allowed to express transgenes for 10-14 days.

Caprola2 recordings in soma of rat hippocampal neurons

*Chemical activation of cultured neurons*. Rat hippocampal neurons seeded in 96-well glass bottom plate were used at 14-16 days *in vitro* (div). Prior to experiments, synaptically blocked neurons were treated with APV/NBQX (25 µM/10 µM) at 37 °C in a humidified 5% $CO_2$ incubator for 30 min. The respective fluorophore and glutamate/glycine (10 µM/2.5 µM) were applied to neuronal cultures at 37 °C in a humidified 5% $CO_2$ incubator for 30 min. Neurons were first washed with 1x HBSS supplemented with HaloTag7 protein (1 µM) for 10 min followed by washing twice with transparent NeuroBasal medium for 10 min each, prior to fixation with 4% PFA and imaging as described above.

*Electric field stimulation of cultured neurons*. Rat hippocampal neurons seeded in 24-well glass bottom plate were used at 14–16 div. Prior to stimulation, a synaptic blocker solution (APV/NBQX (25 µM/10 µM) in NeuroBasal medium) was applied to cells at 37 °C in a humidified 5% $CO_2$ incubator for 30 min and the fluorophore substrate was applied for 10 min. Neuron cultures were placed on a pre-warmed (37 °C) widefield microscope stage in 5% $CO_2$ atmosphere and a custom-build 24-well cap stimulator with platin electrodes linked to a stimulation control unit was mounted on top[16]. In order to elicit defined trains of action potentials, stimulation patterns of 80 Hz, 100 mA and 1 ms pulse width were delivered to live neurons in the presence of the fluorophore substrate: 10 short sequences of stimulation with different numbers of action potentials (1 to 100 AP per sequence) over the course of 6 min, interspaced with 20 s resting periods. After stimulation, cells were washed with 1x PBS supplemented with HaloTag7 protein (1 µM) for 10 min followed by fixation with 4% PFA. Fluorescence imaging was conducted as described above in 1x PBS at 37 °C.

Postsyn-Caprola2 localization and recording in rat hippocampal neurons

*Chemical activation of cultured neurons*. Rat hippocampal neurons seeded in 96-well glass bottom plate were used at 18-20 div. Prior to experiments, synaptically blocked neurons were treated with APV/NBQX (25 µM/10 µM) at 37 °C in a humidified 5% $CO_2$ incubator for 30 min. The fluorophore $JF_{669}$-CA (25 nM) and KCl (10 mM, only in stimulated conditions) were applied to neuronal cultures at 37 °C in a humidified 5% $CO_2$ incubator for 25 min. Neurons were first washed with NeuroBasal medium supplemented with HaloTag7 protein (1 µM) for 5 min followed by one wash with transparent 1x PBS (Gibco) prior to fixation with pre-warmed 4% PFA. Fixed samples were stored at 4 °C in 1x PBS (Gibco) and stained within one week.

*Immunofluorescence staining.* Fixed neurons were permeabilized with 0.3% Triton X-100 in 1x PBS (Gibco) for 3 min at ambient temperature and washed thrice for 10 min with washing buffer (Method Table 1). Samples were incubated with respective antibodies (1:300 dilution in WB buffer of camelid sdAB FluoTag-X4 anti-GFP labeled with ATTO488 (NanoTag Biotechnologies); 1:333 dilution in WB buffer of camelid sdAB FluoTag-X2 anti-PSD95 labeled with AZDye568 (NanoTag Biotechnologies); 1:800 dilution in WB buffer of guinea pig anti-MAP2 polyclonal AB (Synaptic Systems)) for 1 h at ambient temperature. After three washes (3x 10 min) with washing buffer (Method Table 1), guinea pig anti-MAP2 primary antibody was labeled with goat anti-guinea pig IgG (1:400 dilution in WB buffer of H&L Alexa Fluor®750 (Abcam Limited)) for 1 h at ambient temperature. DAPI (Thermo Fisher Scientific) was added to the secondary antibody incubation step at a final concentration of 0.1 µg $mL^{-1}$. Neurons were washed thrice with 1x PBS (Gibco) for 10 min at ambient temperature, followed by storage in 1x PBS (Gibco) at 4 °C and imaging within one week after staining.

*Image acquisition and analysis.* Images were acquired on a Nikon Ti2-E inverted microscope coupled to a Crest X-Light V2/V3 spinning disk confocal module (Nikon). The microscope was equipped with a CFI Plan Apo Lambda D 60× oil objective (NA 1.42, WD 0.15 mm; Nikon), a Celesta light engine (Lumencor), and a Kinetix 10.2 MP back-illuminated sCMOS camera (Photometrics). Images were acquired as z-stacks of up to 75 optical sections with a step size of 0.2 $\mu$m.

For postsynaptic co-localization analysis in primary neurons, maximum projections were generated for the multi-channel images, background was subtracted using the rolling ball algorithm in Fiji, and colocalization between postsyn-Caprola2 and PSD95 immunofluorescence was quantified using the JACoP plugin[17] in Fiji[11].

For postsynaptic Caprola2 activity measurements, multichannel fluorescence images were batch processed using custom Fiji[11] macros. Maximum-intensity projections were generated for all channels and background subtraction was applied. A binary mask to define localization of postsyn-Caprola2 in synaptic compartments was generated for the mGreenLantern signal applying Gaussian blur, intensity thresholding, watershed separation and particle-based filtering to obtain individual synaptic regions of interest (ROIs). The resulting ROIs were applied to the corresponding GreenLantern and $JF_{669}$-CA maximum projections, and the mean fluorescence intensity of both channels was extracted for each synapse. Fluorescence measurements were exported for downstream analysis in R[5].

$JF_{669}$-CA fluorescence was normalized to the corresponding GreenLantern fluorescence on a per-synapse basis to account for differences in reporter expression and subsequently normalized to the median signal of the no-fluorophore control. For statistical analyses, synapse-level measurements were summarized per field of view, which served as the experimental unit.

Statistical analyses and activity-state classification were performed in R (version 2026.01.1+403) using custom scripts. Synapse-level measurements were used for visualization, whereas fields of view were treated as the experimental unit for statistical inference. Pairwise comparisons were performed using two-sided Wilcoxon rank-sum tests with Benjamini–Hochberg correction for multiple testing unless otherwise stated.

Postsyn-Caprola2 recordings in mouse hippocampal neurons

*Chemical activation of cultured neurons.* Mouse hippocampal neurons seeded onto ø18 mm glass coverslips were used at 24 div. Neurons were preincubated in extracellular neuron solution (Method Table 1) supplemented with TTX (1 $\mu$M) or picrotoxin (50 $\mu$M) or without specific blockers for 10 min at 37 °C in a humidified 5% $CO_2$ incubator before addition of 0.5 nM CPY-CA for 90 min. After incubation, neurons were washed once in extracellular neuron solution lacking CPY-CA and imaged in the presence of TTX or picrotoxin.

*Image acquisition.* Images were acquired on an inverted microscope (Nikon Eclipse Ti2) equipped with a 60x/1.45 NA objective and an sCMOS camera (Orca Flash, Hamamatsu Photonics, Japan). Images were acquired using the software NIS elements (Nikon) at a frame rate of 20 Hz for 50 frames.

*Image analysis.* Maximum projections were generated, and background was subtracted using the rolling ball algorithm in Fiji[11]. The mGreenLantern fluorescent signal was used to localize individual synapses along the dendritic tree and normalize the corresponding fluorescent intensity of CPY-CA fluorescent signal. To quantify the activity of individual synapses the ratio between the fluorescent intensity of CPY-CA and mGreenLantern within a circular region of 10 pixel in diameter was calculated. Data were analysed and plotted using GraphPad Prism (version 10) after export from Fiji[11].

### *In vivo* experiments

General remarks on zebrafish husbandry

Adult and larval zebrafish (Danio rerio) were maintained (fish husbandry, permit number 35-9185.64-BH to Jochen Wittbrodt) in accordance with local animal welfare standards (Tierschutzgesetz §11, Abs. 1, Nr. 1) and with European Union animal welfare guidelines. The fish facility is under the supervision of the local representative of the animal welfare agency.

After natural mating, up to 100 embryos were kept in petri dishes with E3 medium, which was exchanged every second day. The fish developmental stage was reported in days post-fertilization (dpf), corresponding to staging at standard temperature of 28.5 ˚C [Kimmel et al 1995]. All embryos used in this work partially carried the Casper or Nacre genetic background, having the mitfa-/-[White et al 2008] and mpv17-/-[D'Agati et al 2017] mutations and melanophore and iridophore pigmentation deficiency for improved optical accessibility[Lister et al 1999]. To remove residual pigmentation, larvae were incubated with 1-phenyl 2-thiourea (PTU, 200 $\mu$M) from 1 dpf until the beginning of the labeling experiment. PTU was exchanged every second day. The herein described experiments on zebrafish larvae, followed by their euthanization, were performed before 5 dpf.

Generation of zebrafish linesFor the generation of transgenic zebrafish lines expressing Caprola11, an animal protocol was approved by the regional government of Oberbayern (Regierung von Oberbayern) with permit number ROB-55.2-2532.Vet_02-21-70 (Herwig Baier). For the generation of transgenic zebrafish lines expressing Caprola21, an animal protocol was approved by the regional government (Regierungspräsidium Tübingen) with permit number 35-9185.81/G-76/24 (Jochen Wittbrodt).

For the generation of transgenic Caprola21 zebrafish, WT AB zebrafish eggs were injected with mRNA encoding the Tol2 transposase [Suster et al 2011] together with a pTol2 plasmid encoding Caprola21 driven by a partial elavl3 pan-neuronal promoter (Tg[elavl3:NES-Caprola21-mEGFP]). At 3 dpf, fluorescent and healthy embryos were sorted under an epifluorescence microscope (excitation 488 nm, emission filter 530/30 nm) and raised to adulthood (F0 generation). F0 individual fish were outcrossed to zebrafish with the Casper genetic background and screened for germ line propagation of the Caprola construct. The offspring with strongest fluorescence were raised to adulthood, constituting the F1 generation. F1 individual fish were outcrossed to zebrafish with the Casper genetic background and screened for germ line propagation of the Caprola constructs. The offspring with strongest fluorescence were raised to adulthood, constituting the F2 generation. Incross offspring (F3) from the F2 generation were used for labeling experiments.

Free swimming labeling experiment

Zebrafish larvae (4 dpf) expressing either Caprola11 or Caprola21 were placed in a 24-well plate containing 500 $\mu$L of E3 medium. Experiments were initiated by adding CPY-CA to reach a final dye concentration of 5 µM. Experiments in both zebrafish lines were conducted in parallel on the same day. Zebrafish larvae were incubated for 0.5–1.5 h in light. To terminate the experiment, larvae were de-stained in E3 medium three times for 20 min and finally 60 min prior to imaging. Zebrafish larvae were maintained in E3 medium until mounting and imaging.

Confocal microscopy of zebrafish larvae

Prior to imaging, zebrafish larvae were mounted on the bottom of glass-bottom 100 mm × 35 mm plastic dishes (4 mL, MatTek, 10 mm glass diameter, uncoated) using 1.3% low melting agarose containing Tricaine (0.16 mg/mL). Confocal microscopy was performed on a commercial Leica Stellaris 5 microscope equipped with a white line laser (WLL), and a hybrid photodetector for single molecule detection (HyD SMD detector). The following settings were used for image acquisition: 20x/0.8 air objective; image size: 775.8 µm x 775.8 µm (1024x1024 pixel); scan speed 600 MHz; pinhole 1 airy unit; zoom 0.75; line averages (6) and 16-bit depth. Z-stacks were performed with 5 µm step sizes. Images were analysed as described above (Imaging data analysis).

### Data representation, reproducibility and statistical analysis

Numerical data was plotted with ggplot2 R packages[5,6]. Imaging data was analyzed and represented using FIJI[11]. Schemes and figures were made with Adobe Illustrator. Biochemical experiments were performed in three technical replicates. Unless stated otherwise, cell experiments were performed in two independent biological replicates including technical replicates. For mammalian cell experiments, biological duplicates are defined as different passages of cell lines. For neurons, biological replicates are defined as different neuron preparations. For zebrafish larvae, individual larvae were considered biological replicates and two independent experiments were conducted at least at two different days from different clutches. Statistical analysis was performed with GraphPad Prism 8.0 or custom R scripts[5,6]. Statistical significances (*i.e.* *p*-values) were calculated using the two-tailed Student's t-test applying Welch's correction or the Kruskal-Wallis/Dunn's test. The different *p*-values were classified as **** for $p$-value ≤ 0.0001, *** for $p$-value ≤ 0.001, ** for $p$-value ≤ 0.01, * for $p$-value ≤ 0.05 and non-significant (n.s.) for $p$-value > 0.05.

**Method Table 1**. Composition of buffers used in the study.

| Buffer | Composition |
|---|---|
| IMAC lysis buffer | 50 mM $KH_2PO_4$, 150 mM NaCl, 5 mM imidazole, 1 mM PMSF, 0.25 mg $mL^{-1}$ lysozyme, pH 8.0 |
| IMAC wash buffer | 50 mM $KH_2PO_4$, 300 mM NaCl, 10 mM imidazole, pH 7.5 |
| IMAC elution buffer | 50 mM $KH_2PO_4$, 300 mM NaCl, 500 mM imidazole, pH 7.5 |
| Activity buffer | 50 mM HEPES, 50 mM NaCl, pH 7.3 |
| FP buffer | 50 mM HEPES, 250 mM NaCl, 0.5 g $L^{-1}$ BSA, 200 $\mu$M EGTA, pH 7.3 |
| Washing buffer | 1x PBS, 2% BSA, 25 mM glycine |
| Extracellular neuron solution | 145 mM NaCl, 2.5 mM KCl, 10mM Glucose, 10mM HEPES, 2 mM $CaCl_2$, pH 7.4 |
| E3 medium | 0.3 g red sea salt in 1 L VE water |

### Supplementary Figures

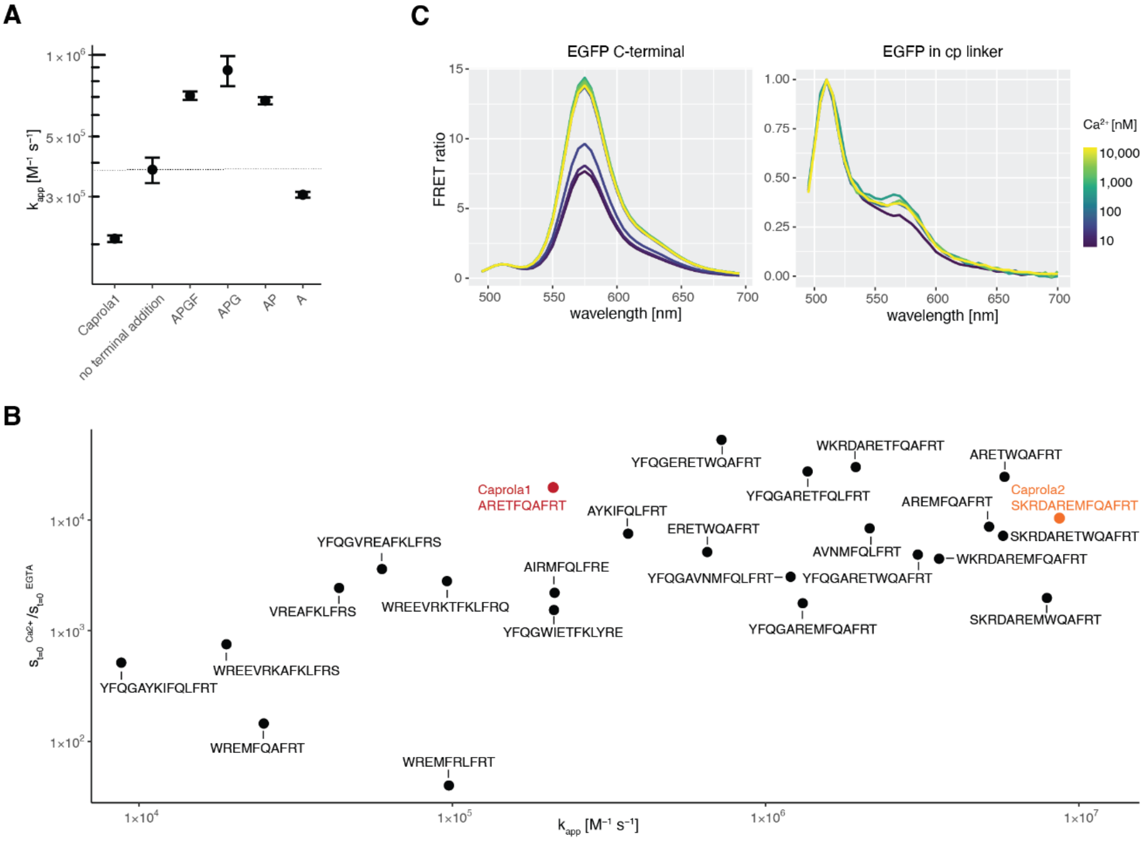

**Figure S1.** Engineering of Caprola2 and Caprola3. (A) Effect of C-terminal extension at residue 141 of cpHaloΔ on labeling kinetics in presence of saturating $Ca^{2+}$. Caprola1 featured an N-terminal Strep-tag connected via enterokinase cleavage site and a C-terminal His-tag connected via linker with residues APGFSSISA. Other constructs featured an N-terminal His-Tag connected via TEV cleavage site, additional point mutation [A151L] in Hpep1, and various C-terminal extensions as indicated. (B) Hpep screening in Caprola3 context (C-terminal residues AP and remodeled cp linker 1, see Figure S5). The ratio of initial slopes of labeling kinetics in presence or absence of $Ca^{2+}$ is plotted against the apparent second order rate constant ($k_{app}$) of labeling reaction in presence of $Ca^{2+}$. (C) $Ca^{2+}$-dependency of FRET of TMR-labeled Caprola2 variant with mEGFP fused to C-terminus (left, intermediate Caprola variant with Hpep YFQGARETFQAFRT and terminal residues AP, cp linker 1 modeled by Rosetta for Caprola3, see Figure S5) or positioned in cp-linker (right, intermediate Caprola variant with Hpep YFQGARETFQAFRT and terminal residues AP).

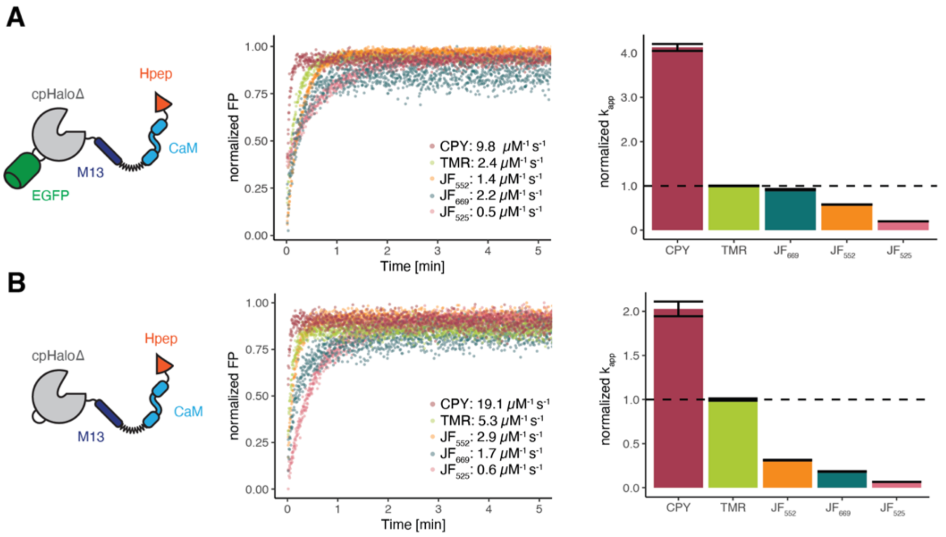

**Figure S2.** Labeling reaction of (A) Caprola2$_9$ and (B) Caprola3$_9$ with different fluorophore substrates. Left: Construct scheme. Middle: Labeling kinetics of protein (50 nM) in the presence of $Ca^{2+}$ (5 mM $CaCl_2$) and apparent second-order rate constants with the different fluorophore substrates (10 nM). Fluorescence polarization values were normalized for each fluorophore to its fully bound fluorescence polarization (normalized FP). Right: Apparent second-order rate constants ($k_{app}$) normalized to the $k_{app}$ with TMR-CA. HaloTag substrates used: CPY-CA, TMR-CA, JF$_{669}$-CA, JF$_{552}$-CA and JF$_{525}$-CA.

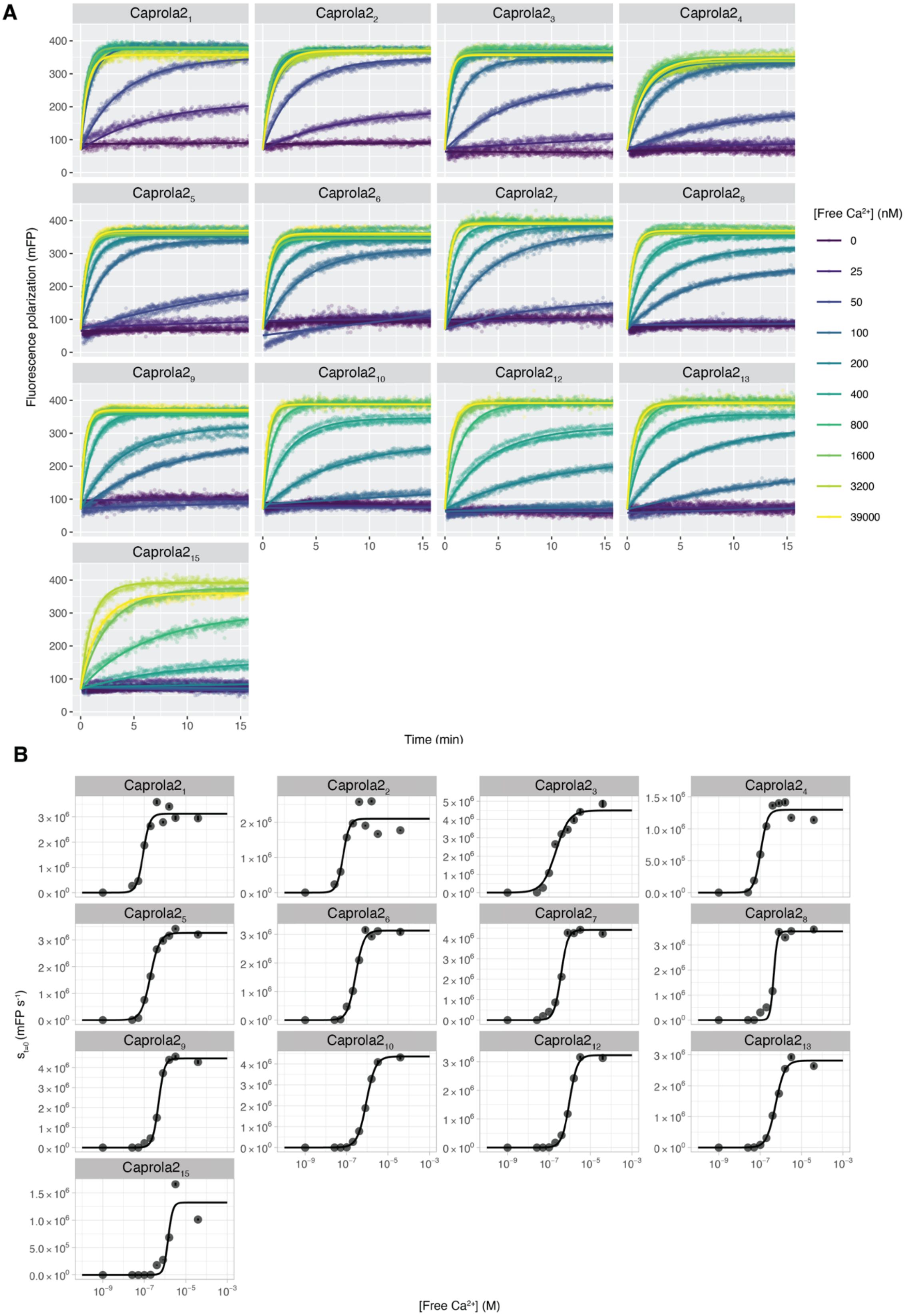

**Figure S3.** Calcium sensitivity of Caprola2 variants. (A) Labeling kinetics of Caprola2 variants (10 nM) with TMR-CA (2 nM) at different concentrations of free $Ca^{2+}$ were followed by fluorescence polarization. A second-order reaction model or a linear model for reactions not reaching a plateau was fit to the data and initial reaction rates ($s_{t=0}$) were determined. (B) Initial reaction rates ($s_{t=0}$) were plotted against free $Ca^{2+}$ concentration and a sigmoidal model was fit to the data to estimate $EC_{50}$ values.

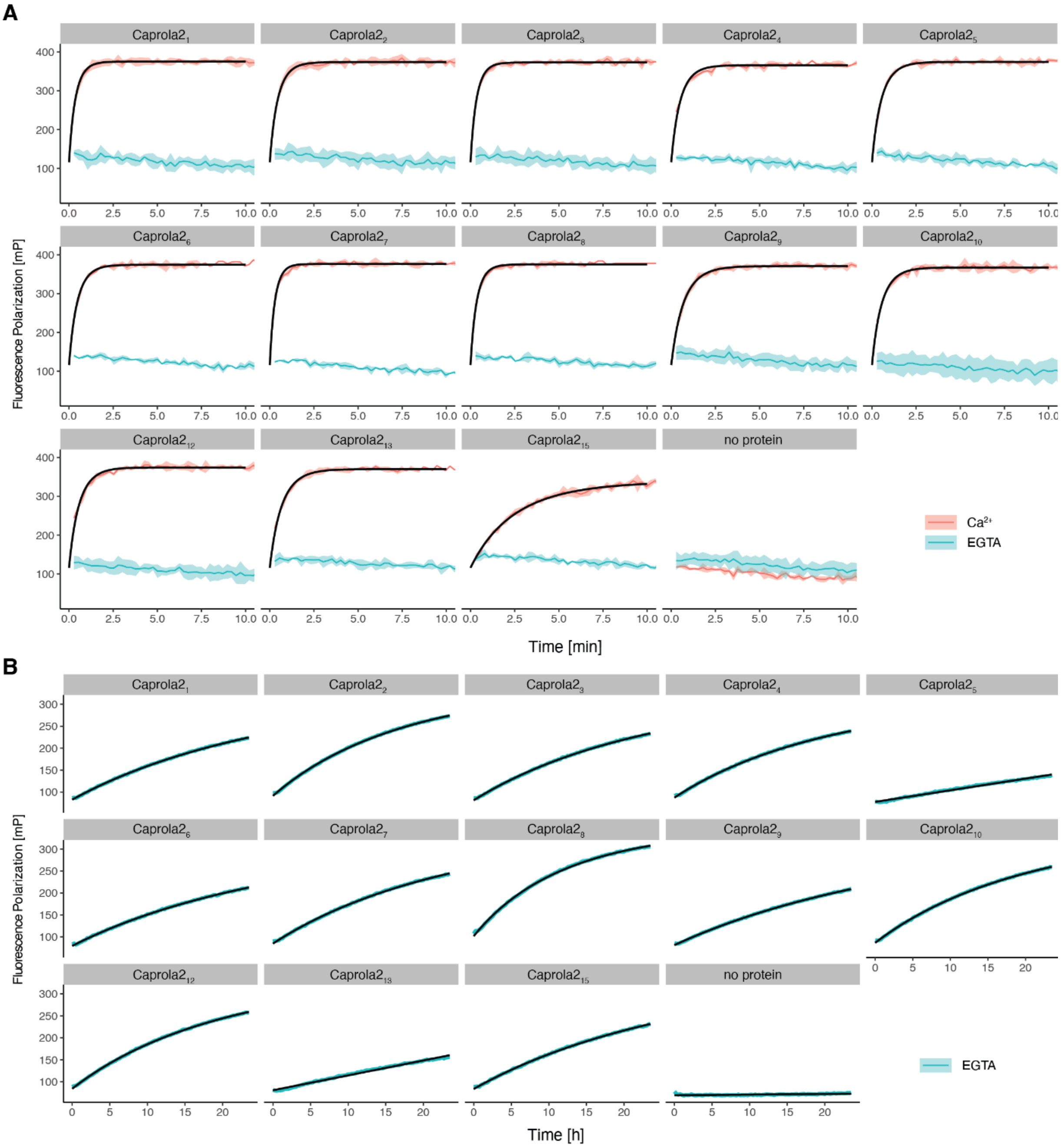

**Figure S4.** Labeling kinetics of the different Caprola2 variants. (A) Labeling kinetics of Caprola2 variants (10 nM) with TMR-CA (2 nM) in presence or absence of free $Ca^{2+}$ measured by fluorescence polarization. A second-order reaction model was fitted to the data in presence of $Ca^{2+}$ to determine apparent second order rate constants (see Table S2). (B) Rate constants in the absence of $Ca^{2+}$ were determined in a long term (24 h) fluorescence polarization assay. A second-order reaction model (with fixed plateau) was fitted to the data to determine apparent second order rate constants (see Table S2).

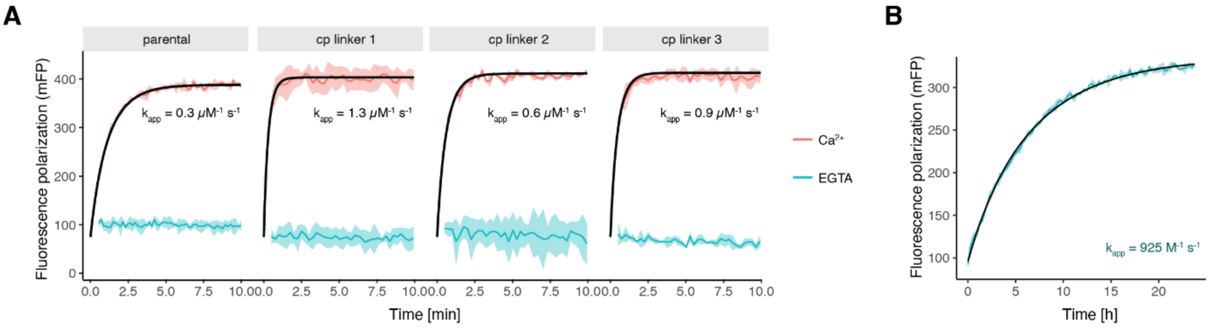

**Figure S5.** Engineering of Caprola3. (A) Labeling kinetics of CaprolaX variants with different cp linkers (50 nM) with TMR-CA (10 nM) in presence of saturating $Ca^{2+}$ and absence of $Ca^{2+}$ (EGTA). Parental: Caprola3 but with Hpep YFQGARETFQAFRT. (B) Labeling kinetics and apparent second order rate constant ($k_{app}^{EGTA}$) of final Caprola3 (50 nM) with TMR-CA (10 nM) in absence of $Ca^{2+}$ (EGTA) over 24h.

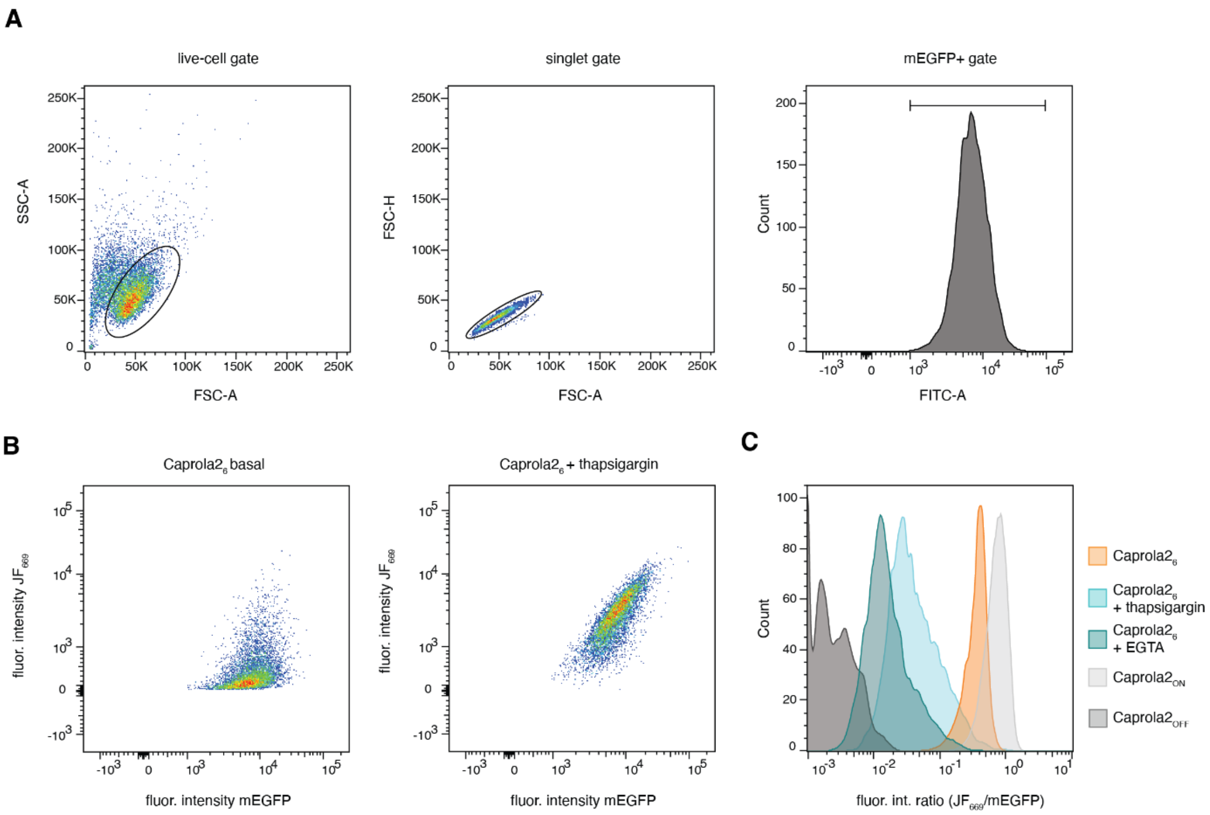

**Figure S6.** Exemplary flow cytometry gating and analysis strategy for U2OS cells expressing Caprola2. (A) Gating strategy of U2OS cells expressing Caprola2$_6$: live-cells (FSC-A x SSC-A), singlets (FSC-A x FSC-H) and mEGFP positive cells (mEGFP+, fluor. int. of mEGFP > $10^3$) are gated. (B) Fluorescence intensities (mEGFP x $JF_{669}$-CA) of U2OS cells expressing Caprola2$_6$ incubated with 10 nM $JF_{669}$-CA for 30 min in the absence (basal) or presence of thapsigargin (100 nM). (C) Histogram of fluorescence intensity ratios ($JF_{669}$-CA /mEGFP) from U2OS cells expressing Caprola2$_6$, Caprola2$_{ON}$ or Caprola2$_{OFF}$ incubated with 10 nM $JF_{669}$-CA for 30 min in the presence or absence of thapsigargin (100 nM) or EGTA (1 mM).

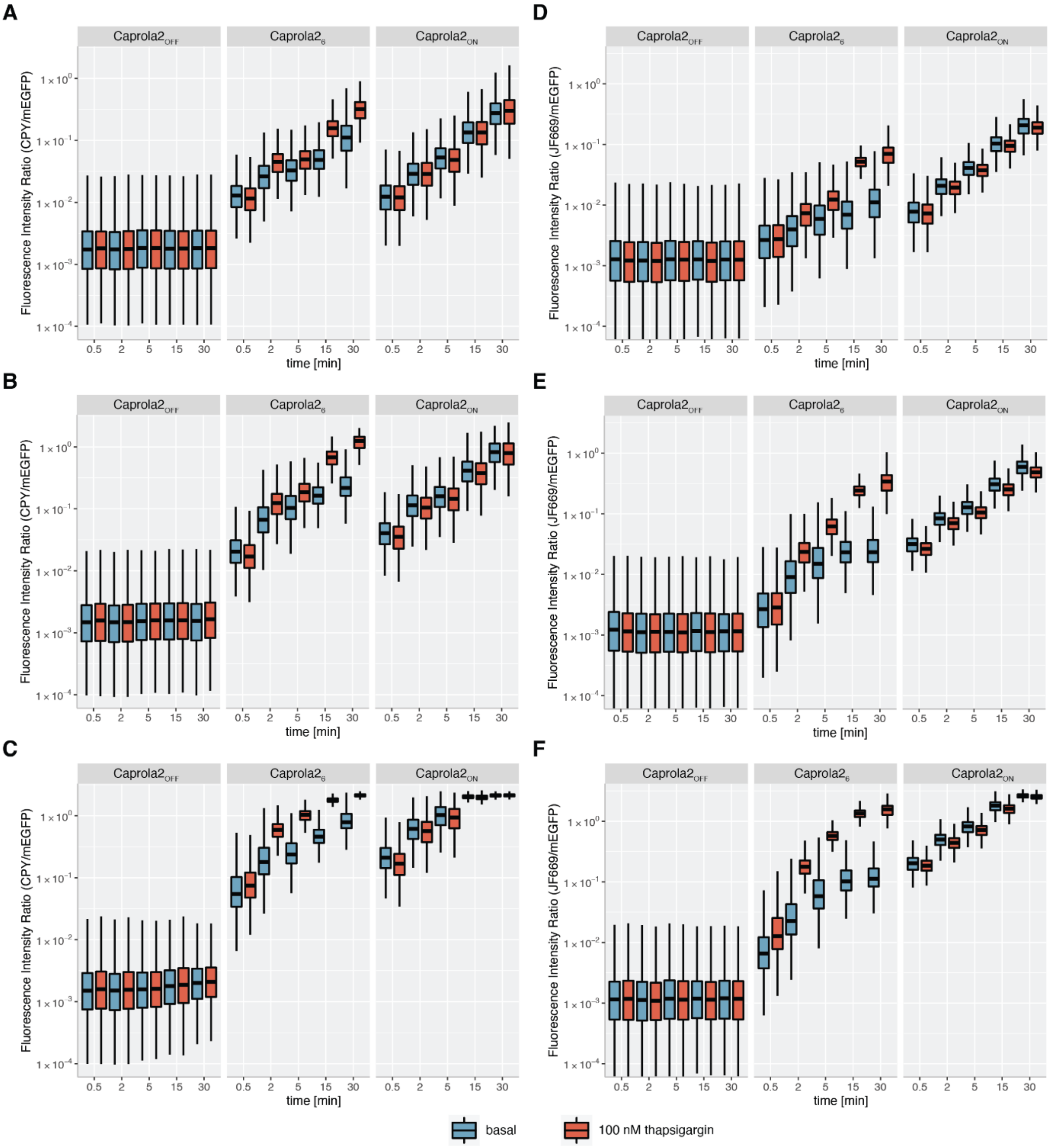

**Figure S7.** Time and fluorophore substrate concentration dependency of Caprola2 labeling in U2OS cells. (A, B, C) Fluorescence intensity ratios (CPY-CA/mEGFP) of U2OS cells expressing Caprola$2_6$, Caprola$2_{OFF}$ and Caprola$2_{ON}$ incubated with either (A) 2 nM, (B) 10 nM or (C) 50 nM CPY-CA and increasing incubation times in the presence or absence of 100 nM thapsigargin are shown as box-and-whisker plots. (D, E, F) Fluorescence intensity ratios ($JF_{669}$-CA/mEGFP) of U2OS cells expressing Caprola$2_6$, Caprola$2_{OFF}$ and Caprola$2_{ON}$ incubated with either (D) 2 nM, (E) 10 nM or (F) 50 nM $JF_{669}$-CA and increasing incubation times in the presence or absence thapsigargin (100 nM) are shown as box-and-whisker plots.

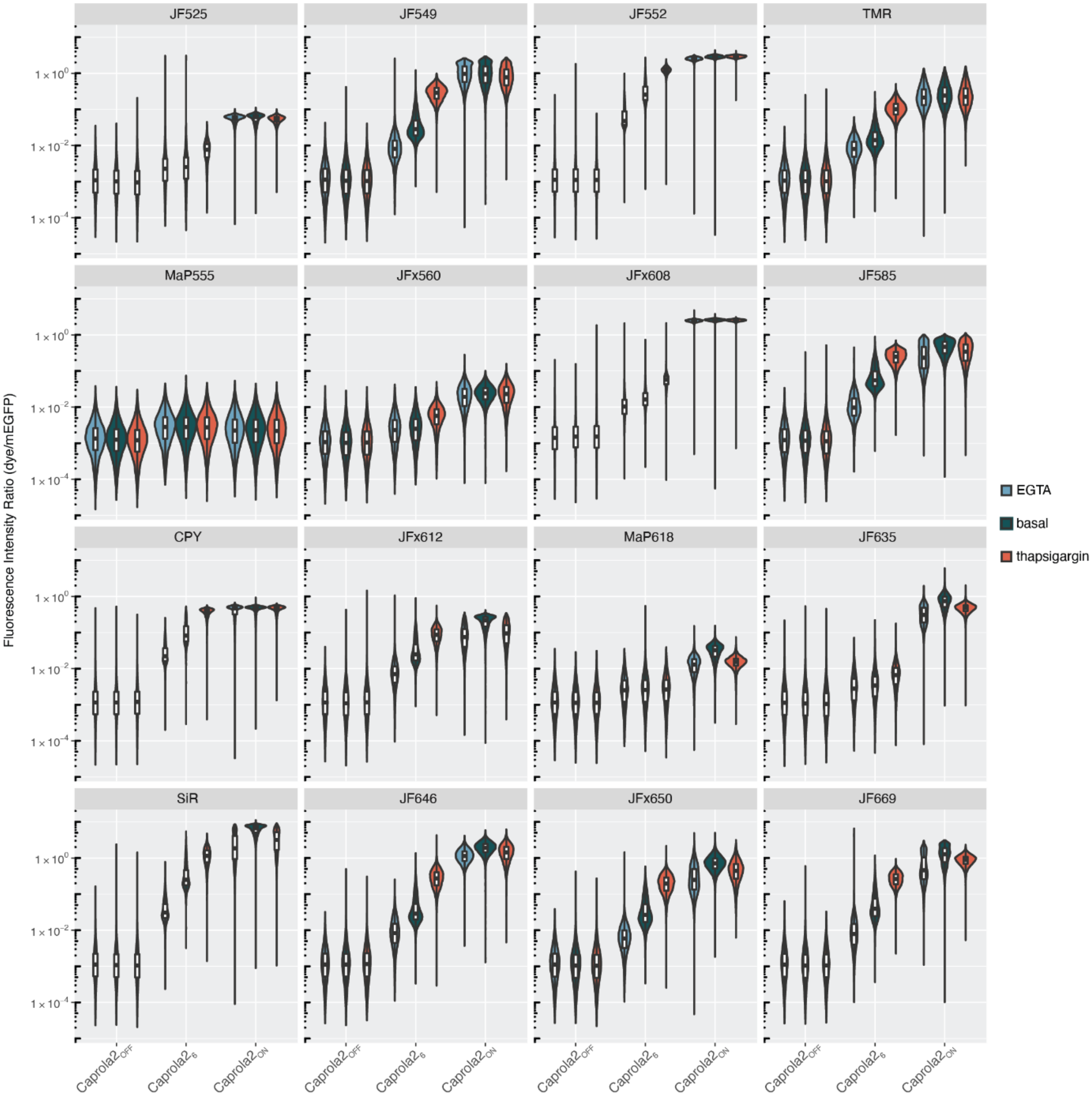

**Figure S8.** Labeling of Caprola2 in U2OS cells with different dyes. (A) Flow cytometry analysis of U2OS cells expressing Caprola2$_6$, Caprola2$_{OFF}$ or Caprola2$_{ON}$ incubated with 10 nM dye for 30 min in the presence of 100 nM thapsigargin or 1 mM EGTA or in the absence of stimulus (basal). Violin plots and box-and-whisker plots of fluorescence intensity ratio (dye/mEGFP) are displayed.

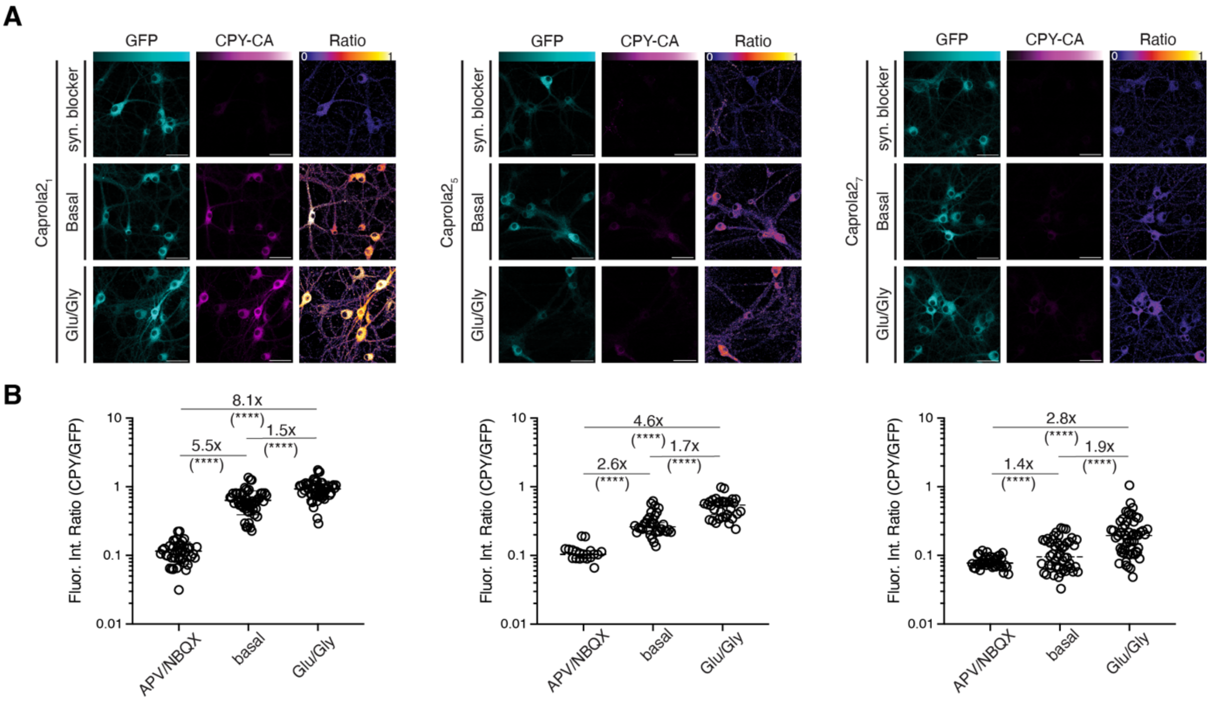

**Figure S9.** Effect of calcium sensitivity of Caprola2 variant on signal intensity in drug-stimulated primary rat hippocampal neurons. (A) Fluorescence microscopy images of Caprola2$_1$, Caprola2$_5$, and Caprola2$_7$ expressing primary rat hippocampal neurons (14 div) after incubation with 10 nM CPY-CA for 30 min without stimulation (basal), or in the presence of either synaptic blockers (30 min preincubation) or glutamate/glycine. (B) Quantification of experiment described in (A) ($N > 20$ cells, ****: $p < 0.0001$; Welch's t test). The center line indicates the median, and error bars indicate 25% and 75% quantiles. Scale bars: 20 $\mu$m.

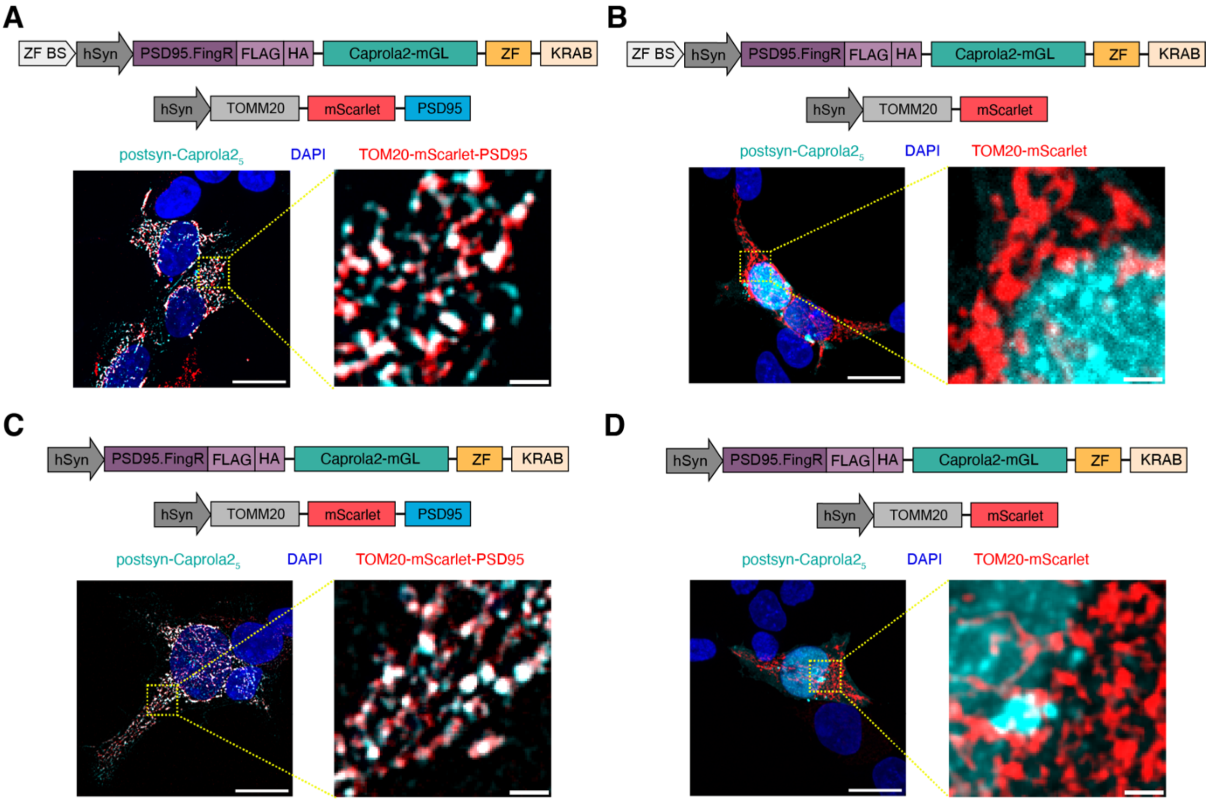

**Figure S10.** Colocalization of postsyn-Caprola2$_5$ and TOM20-PSD95 in HEK293T cells. Scheme of the composition of postsyn-Caprola2$_5$ and target/non-target constructs on top and exemplary images of their co-expression in HEK293T cells on bottom. (A) Exemplary image of postsyn-Caprola2$_5$ with ZF binding site upstream of hSyn promoter, co-expressed with TOM20-mScarlet-PSD95. (B) Exemplary image of postsyn-Caprola2$_5$ with ZF binding site upstream of hSyn promoter, co-expressed with TOM20-mScarlet lacking target. (C) Exemplary image of postsyn-Caprola2$_5$ without ZF binding site upstream of hSyn promoter, co-expressed with TOM20-mScarlet-PSD95. (D) Exemplary image of postsyn-Caprola2$_5$ without ZF binding site upstream of hSyn promoter, co-expressed with TOM20-mScarlet lacking target. Scale bars: overview image: 20 µm; zoom: 2 µm.

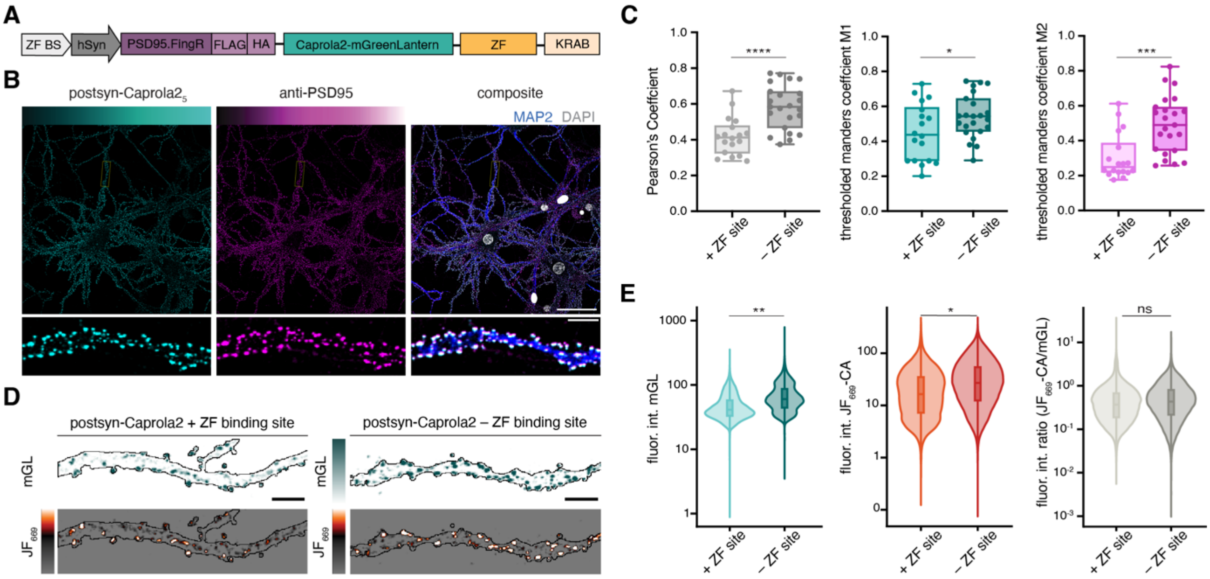

**Figure S11.** Evaluation of postsyn-Caprola2$_5$ expression feedback loop. (A) Scheme of postsyn-Caprola2$_5$ with CCR5-L zinc finger binding site (ZF BS) upstream of the hSyn promoter. (B) Representative overview (top) fluorescence images and dendritic sections (bottom) of primary rat hippocampal neurons expressing postsyn-Caprola2$_5$ with CCR5-L zinc finger binding site (left) and immunofluorescence staining of PSD95 (middle). Their merge (right) shows areas of colocalization in white, the dendritic marker MAP2 in blue and DAPI in grey. Scale bars: full image: 50 $\mu$m, dendritic section: 5 $\mu$m. (C) Colocalization analysis of postsyn-Caprola2$_5$ mGreenLantern signal and anti-PSD95 signal from experiment described in (B) and in Figure 3C. Pearson's coefficient, and thresholded Mander's coefficients M1 and M2 (unpaired two-tailed t-test, *: $p \leq 0.05$, ***: $p \leq 0.001$, ****: $p \leq 0.0001$). The variant without ZF binding site showed significantly improved co-localization with PSD95, targeting specificity (M1), and synaptic saturation (M2). (D) Exemplary fluorescence images of dendritic sections of primary rat hippocampal neurons expressing either postsyn-Caprola2$_5$ with CCRL-5 zinc finger binding site upstream of promoter (left) or without CCRL-5 binding site, labeled with $JF_{669}$-CA (25 nM, 25 min). Scale bars: 5 $\mu$m. (E) Quantification of postsyn-Caprola2$_5$ labeling from experiment described in (D) (unpaired two-tailed t-test, ns: $p > 0.05$, *: $p \leq 0.05$, **: $p \leq 0.01$). Comparison of mGreenLantern fluorescence intensity (mGL, left), $JF_{669}$-CA fluorescence intensity (middle) and fluorescence intensity ratios (mGL/$JF_{669}$-CA, right).

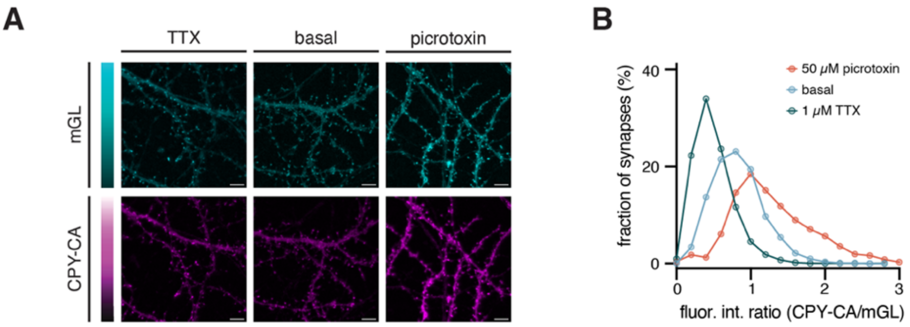

**Figure S12.** Recording post-synaptic activity profiles over prolonged time period of 90 min. (A) Fluorescence microscopy images of primary mouse hippocampal neurons expressing postsyn-Caprola$2_5$ labeled with CPY-CA (0.5 nM, 90 min) in the presence of synaptic blockers TTX (1 $\mu$M) or picrotoxin (50 $\mu$M) or without treatment in medium lacking $Mg^{2+}$. Scale bars: 5 $\mu$m. (B) Quantification of fluorescence intensity ratios (CPY-CA/mGreenLantern) of postsyn-Caprola$2_5$ labeling from experiment described in (A) (N $\geq$ 2,795 synapses, histogram displaying mean ratio ± s.d.).

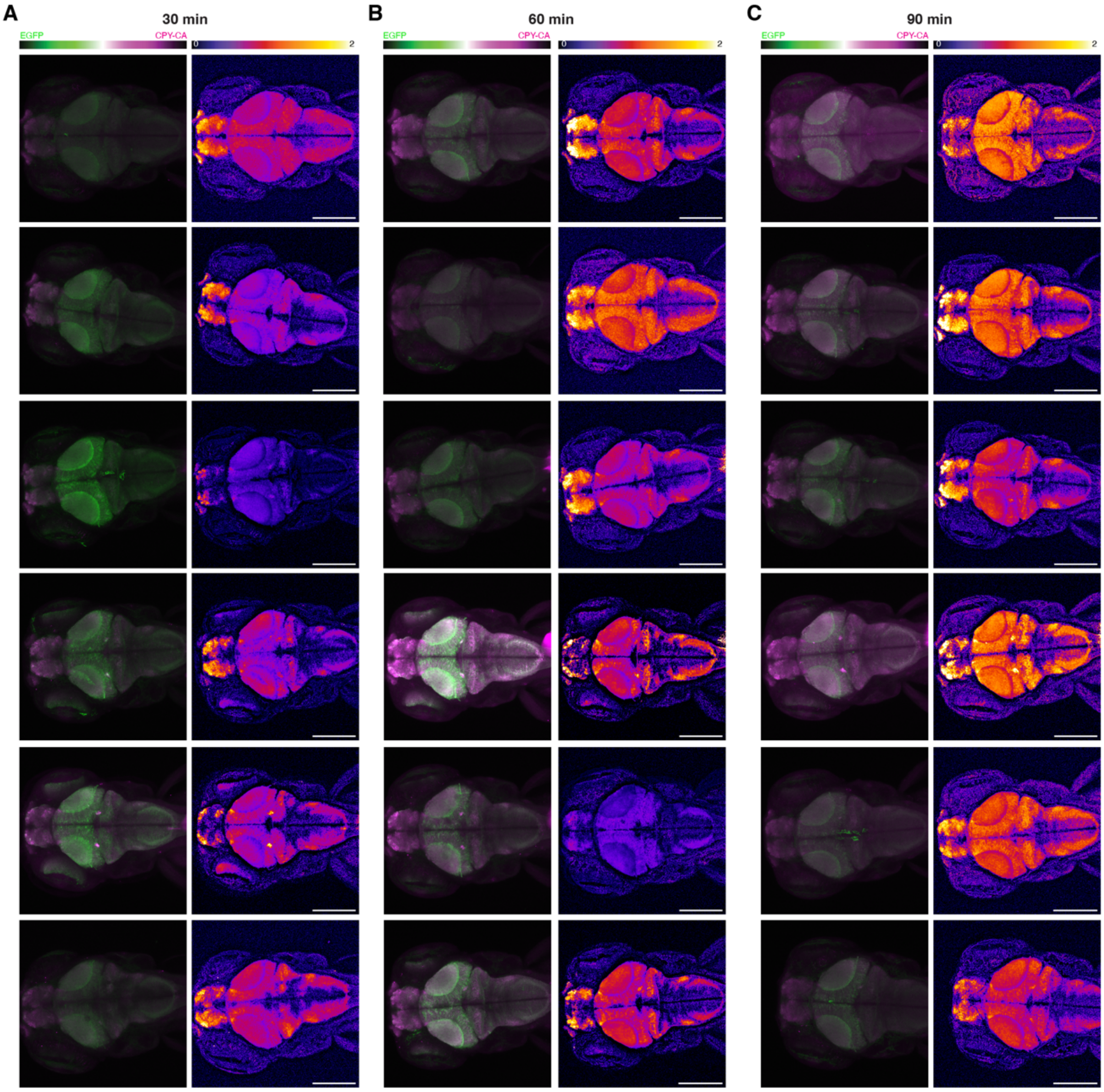

**Figure S13.** Caprola2$_1$ labeling in the zebrafish larval brain (4 dpf). Maximum intensity projections of freely-swimming zebrafish larvae incubated with CPY-CA (5 $\mu$M) for the indicated time interval. Ratio: CPY-CA/mEGFP. Scale bars: 200 $\mu$m.

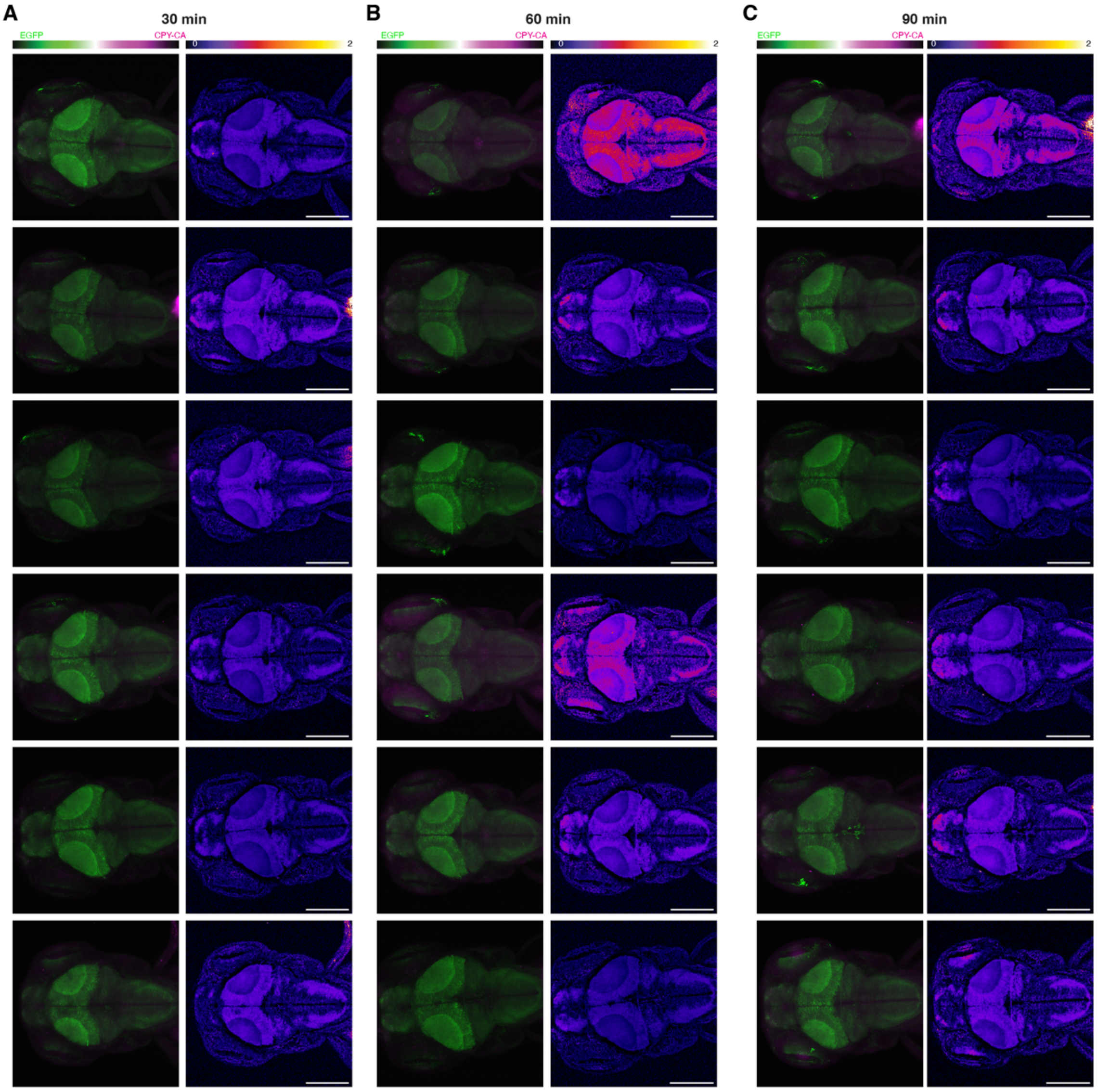

**Figure S14.** Caprola1$_1$ labeling in the zebrafish larval brain (4 dpf). Maximum intensity projections of freely-swimming zebrafish larvae incubated with CPY-CA (5 $\mu$M) for the indicated time interval. Ratio: CPY-CA/mEGFP. Scale bars: 200 $\mu$m.

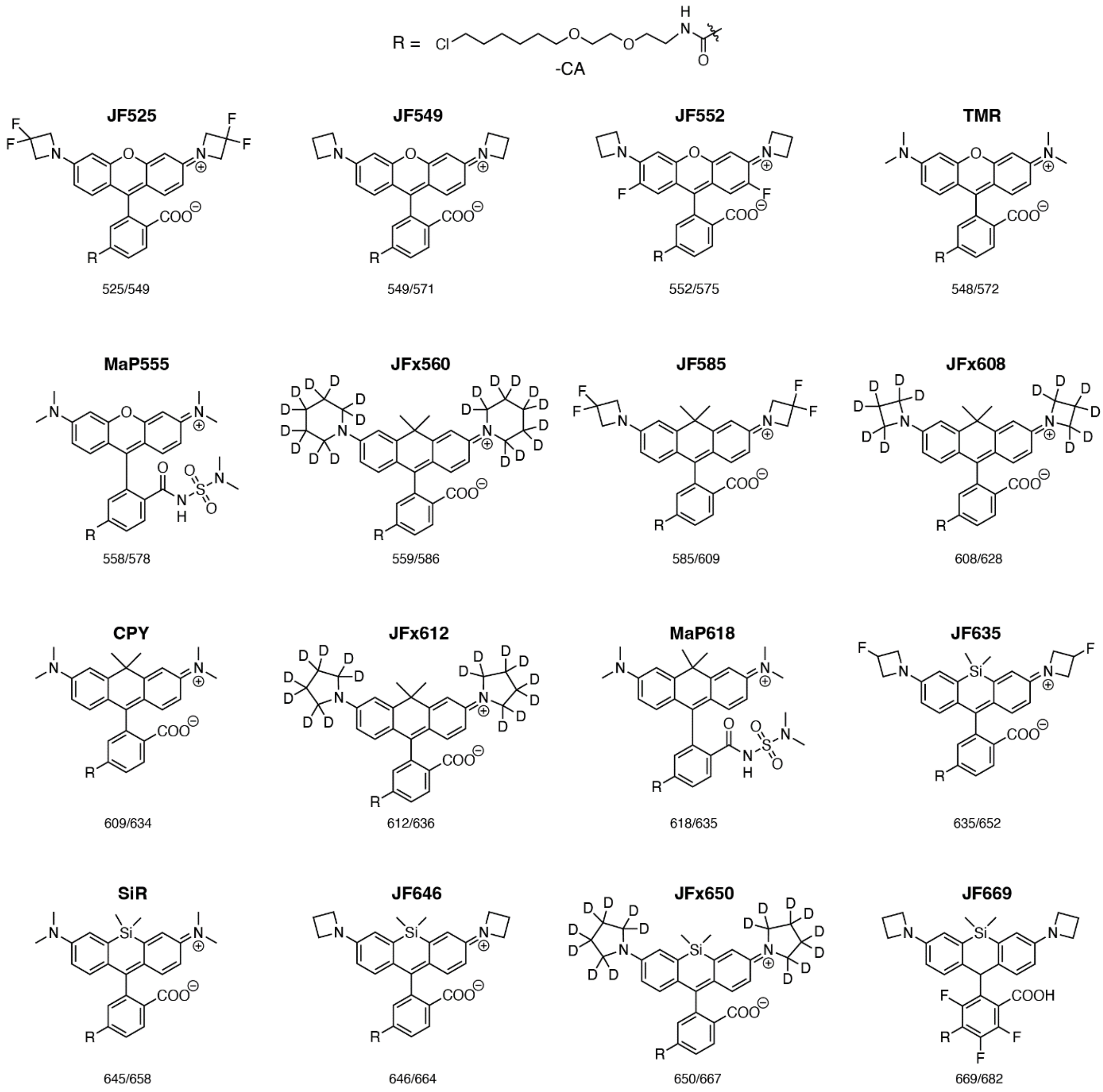

**Figure S15.** Chemical structures of fluorophore HaloTag ligands used in this study. Excitation and emission maxima are given below each structure. Janelia Fluor (JF) fluorescent substrates were kind gifts of L. D. Lavis (Janelia Research Campus, Ashburn, VA, USA). TMR-CA, CPY-CA, SiR-CA, MaP555-CA and MaP618-CA were synthesized in-house.

### Supplementary Tables

**Table S1.** Melting temperatures of selected Caprola variants, determined by nanoDSF in the presence of $Ca^{2+}$ (5 mM $CaCl_2$) or absence of $Ca^{2+}$ (500 $\mu$M EGTA). CaprolaX: intermediate Caprola3 but with Hpep YFQGARETFQAFRT.

| | + $Ca^{2+}$ (°C) | + EGTA (°C) |
|---|---|---|
| Caprola1 | 31.0 | 38.4 |
| Caprola1 incl. mEGFP | 29.3 | 35.6 |
| Caprola2 | 33.4 | 38.2 |
| CaprolaX – $(GGS/T)_5$ linker | 31.1 | 38.0 |
| CaprolaX – cp linker 1 | 33.7 | 40.3 |
| CaprolaX – cp linker 2 | 33.1 | 39.8 |
| CaprolaX – cp linker 3 | 33.3 | 39.7 |
| Caprola3 | 35.0 | 41.0 |

**Table S2.** Biochemical characterization of Caprola2 variants *in vitro.*

| Caprola2 variant | $EC_{50}$ (nM)[a] | $k_{app}^{Ca2+}$ ($M^{-1}s^{-1}$) (± s.d.)[b] | $k_{app}^{EGTA}$ ($M^{-1}s^{-1}$) (± s.d.)[b] | $k_{app}^{Ca2+}/k_{app}^{EGTA}$ (± s.d.)[b] | Hill coefficient [a] |
|---|---|---|---|---|---|
| $Caprola2_1$ | 87(67 - 113) | $(3.94 \pm 0.13) \times 10^6$ | $278.5 \pm 5.4$ | $5043 \pm 241$ | 2.99(1.63 - 5.83) |
| $Caprola2_2$ | 70(47 - 106) | $(4.96 \pm 0.16) \times 10^6$ | $440.6 \pm 7.4$ | $2421 \pm 94$ | 3.17(1.07 - 7.48) |
| $Caprola2_3$ | 207(155 - 277) | $(5.26 \pm 0.15) \times 10^6$ | $284.7 \pm 5.1$ | $4704 \pm 203$ | 1.40(0.96 - 2.14) |
| $Caprola2_4$ | 114(90 - 144) | $(3.52 \pm 0.10) \times 10^6$ | $374.0 \pm 8.6$ | $2831 \pm 145$ | 2.83(1.61 - 5.17) |
| $Caprola2_5$ | 201(180 - 224) | $(3.32 \pm 0.07) \times 10^6$ | $76.7 \pm 0.2$ | $11679 \pm 269$ | 1.98(1.67 - 2.34) |
| $Caprola2_6$ | 271(239 - 308) | $(3.81 \pm 0.10) \times 10^6$ | $241.8 \pm 6.3$ | $4239 \pm 244$ | 2.26(1.74 - 2.95) |
| $Caprola2_7$ | 381(331 - 440) | $(5.09 \pm 0.14) \times 10^6$ | $310.5 \pm 6.4$ | $5271 \pm 258$ | 2.71(1.88 - 3.91) |
| $Caprola2_8$ | 448(382 - 525) | $(6.17 \pm 0.20) \times 10^6$ | $648.3 \pm 11.7$ | $2131 \pm 74$ | 4.93(2.77 - 8.60) |
| $Caprola2_9$ | 500(477 - 524) | $(2.55 \pm 0.05) \times 10^6$ | $237.1 \pm 5.7$ | $3041 \pm 166$ | 3.06(2.50 - 3.80) |
| $Caprola2_{10}$ | 902(861 - 946) | $(3.22 \pm 0.10) \times 10^6$ | $361.3 \pm 5.6$ | $2243 \pm 84$ | 1.95(1.84 - 2.09) |
| $Caprola2_{12}$ | 993(937 - 1052) | $(2.63 \pm 0.05) \times 10^6$ | $347.6 \pm 5.0$ | $2217 \pm 78$ | 2.34(2.02 - 2.73) |
| $Caprola2_{13}$ | 588(513 - 674) | $(3.13 \pm 0.09) \times 10^6$ | $79.5 \pm 0.3$ | $7087 \pm 153$ | 2.03(1.58 - 2.60) |
| $Caprola2_{15}$ | 1200(1177 - 1223) | $(7.00 \pm 0.20) \times 10^5$ | $272.6 \pm 5.7$ | $591 \pm 28$ | 3.82(1.36 - 8.63) |

Biochemical characterization of Caprola2. $EC_{50}$: free $Ca^{2+}$ concentration leading to half maximal Caprola2 labeling rate. $k_{app}^{Ca2+}$: second-order rate constant in saturating $Ca^{2+}$ concentration. $k_{app}^{EGTA}$: second-order rate constant in absence of free $Ca^{2+}$. Hill coefficient: hill coefficient of the sigmoidal model fitted to free $Ca^{2+}$ titrations. [a]95% confidence intervals are given in bracket. [b] s.d.: standard deviation

### Protein sequences

>Caprola1$_9$
(M)ARETFQAFRTGSDQLTEEQIAEFKEAFSLFDKDGDGTITTKELGTVMRSLGQNPTEAEL
QDMINEVDADGDGTIDFPEFLTMMARKMKDTDSEEEIREAFRVFDKDGNGYISAAELRHVM
TNLGEKLTDEEVDEMIREADIDGDGQVNYEEFVVMMTAKEFPPPPPPPPPPPPPPPPPPPP
PPPPPPPPPPPGGSRVDSSRRKMNKTGHALRAIGRLSSLEGGSDVGRKLIIDQNVFIEGTLP
MGVVRPLTEVEMDHYREPFLNPVDREPLWRFPNELPIAGEPANIVALVEEYMDWLHQSPV
PKLLFWGTPGVLIPPAEAARLAKSLPNCKAVDIGPGLNLLQEDNPDLIGSEIARWLSTLEIGG
TGGSGGTGGSGGSIGTGFPFDPHYVEVLGERMHYVDVGPRDGTPVLFLHGNPTSSYVWR
NIIPHVAPTHRCIAPDLIGMGKSDKPDLGYFFDDHVRFMDAFIEALGLEEVVLVIHDWGSALG
FHWAKRNPERVKGIAFMEFIRPIPTWDEW

Hpep; Calmodulin; M13; cpHaloΔ; linker

>Caprola1$_9$ with mEGFP
(M)ARETFQAFRTGSDQLTEEQIAEFKEAFSLFDKDGDGTITTKELGTVMRSLGQNPTEAEL
QDMINEVDADGDGTIDFPEFLTMMARKMKDTDSEEEIREAFRVFDKDGNGYISAAELRHVM
TNLGEKLTDEEVDEMIREADIDGDGQVNYEEFVVMMTAKEFPPPPPPPPPPPPPPPPPPPP
PPPPPPPPPPPGGSRVDSSRRKMNKTGHALRAIGRLSSLEGGSDVGRKLIIDQNVFIEGTLP
MGVVRPLTEVEMDHYREPFLNPVDREPLWRFPNELPIAGEPANIVALVEEYMDWLHQSPV
PKLLFWGTPGVLIPPAEAARLAKSLPNCKAVDIGPGLNLLQEDNPDLIGSEIARWLSTLEIGG
TGGSGGTGGSGGSIGTGFPFDPHYVEVLGERMHYVDVGPRDGTPVLFLHGNPTSSYVWR
NIIPHVAPTHRCIAPDLIGMGKSDKPDLGYFFDDHVRFMDAFIEALGLEEVVLVIHDWGSALG
FHWAKRNPERVKGIAFMEFIRPIPTWDEWGGSMVSKGEELFTGVVPILVELDGDVNGHKFS
VSGEGEGDATYGKLTLKFICTTGKLPVPWPTLVTTLTYGVQCFSRYPDHMKQHDFFKSAMP
EGYVQERTIFFKDDGNYKTRAEVKFEGDTLVNRIELKGIDFKEDGNILGHKLEYNYNSHNVYI
MADKQKNGIKVNFKIRHNIEDGSVQLADHYQQNTPIGDGPVLLPDNHYLSTQSKLSKDPNE
KRDHMVLLEFVTAAGITLGMDELYK

Hpep; Calmodulin; M13; cpHaloΔ; linker; mEGFP

>Caprola2$_9$
(M)SKRDAREMFQAFRTGSDQLTEEQIAEFKEAFSLFDKDGDGTITTKELGTVMRSLGQNPT
EAELQDMINEVDADGDGTIDFPEFLTMMARKMKDTDSEEEIREAFRVFDKDGNGYISAAEL
RHVMTNLGEKLTDEEVDEMIREADIDGDGQVNYEEFVVMMTAKEFPPPPPPPPPPPPPPP
PPPPPPPPPPPPPPPPGGSRVDSSRRKMNKTGHALRAIGRLSSLEGGSDVGRKLIIDQNVFIE
GTLPMGVVRPLTEVEMDHYREPFLNPVDREPLWRFPNELPIAGEPANIVALVEEYMDWLH
QSPVPKLLFWGTPGVLIPPAEAARLAKSLPNCKAVDIGPGLNLLQEDNPDLIGSEIARWLSTL
EIGSGMVSKGEELFTGVVPILVELDGDVNGHKFSVSGEGEGDATYGKLTLKFICTTGKLPVP
WPTLVTTLTYGVQCFSRYPDHMKQHDFFKSAMPEGYVQERTIFFKDDGNYKTRAEVKFEG
DTLVNRIELKGIDFKEDGNILGHKLEYNYNSHNVYIMADKQKNGIKVNFKIRHNIEDGSVQLA
DHYQQNTPIGDGPVLLPDNHYLSTQSKLSKDPNEKRDHMVLLEFVTAAGITLGMDELYKGS
GIGTGFPFDPHYVEVLGERMHYVDVGPRDGTPVLFLHGNPTSSYVWRNIIPHVAPTHRCIA
PDLIGMGKSDKPDLGYFFDDHVRFMDAFIEALGLEEVVLVIHDWGSALGFHWAKRNPERVK
GIAFMEFIRPIPTWDEWAP

Hpep; Calmodulin; M13; cpHaloΔ; linker; mEGFP

>Caprola3₉
(M)SKRDAREMFQAFRTGSDQLTEEQIAEFKEAFSLFDKDGDGTITTKELGTVMRSLGQNPT
EAELQDMINEVDADGDGTIDFPEFLTMMARKMKDTDSEEEIREAFRVFDKDGNGYISAAEL
RHVMTNLGEKLTDEEVDEMIREADIDGDGQVNYEEFVVMMTAKEFPPPPPPPPPPPPPPPP
PPPPPPPPPPPPPPPPGGSRVDSSRRKMNKTGHALRAIGRLSSLEGGSDVGRKLIIDQNVFIE
GTLPMGVVRPLTEVEMDHYREPFLNPVDREPLWRFPNELPIAGEPANIVALVEEYMDWLH
QSPVPKLLFWGTPGVLIPPAEAARLAKSLPNCKAVDIGPGLNLLQEDNPDLIGSEIARWLSTL
EIGTDPREGKTAADDARDKSQASGHIGTGFPFDPHYVEVLGERMHYVDVGPRDGTPVLFL
HGNPTSSYVWRNIIPHVAPTHRCIAPDLIGMGKSDKPDLGYFFDDHVRFMDAFIEALGLEEV
VLVIHDWGSALGFHWAKRNPERVKGIAFMEFIRPIPTWDEWAP

Hpep; Calmodulin; M13; cpHaloΔ; linker

<u>cp linker sequences</u>

cp linker 1 GTDPREGKTAADDARDKSQASGH
cp linker 2 KGAEDAKERLERARDELSKKD
cp linker 3 NIDERQQDWLDKWLKSYNSSTGQ

>Caprola2$_{OFF}$
(M)DQLTEEQIAEFKEAFSLFDKDGDGTITTKELGTVMRSLGQNPTEAELQDMINEVDADGD
GTIDFPEFLTMMARKMKDTDSEEEIREAFRVFDKDGNGYISAAELRHVMTNLGEKLTDEEV
DEMIREADIDGDGQVNYEEFVVMMTAKEFPPPPPPPPPPPPPPPPPPPPPPPPPPPPPPPG
GSRVDSSRRKFNKTAHALRAIGRLSSLEGGSDVGRKLIIDQNVFIEGTLPMGVVRPLTEVEM
DHYREPFLNPVDREPLWRFPNELPIAGEPANIVALVEEYMDWLHQSPVPKLLFWGTPGVLI
PPAEAARLAKSLPNCKAVDIGPGLNLLQEDNPDLIGSEIARWLSTLEIGSGMVSKGEELFTG
VVPILVELDGDVNGHKFSVSGEGEGDATYGKLTLKFICTTGKLPVPWPTLVTTLTYGVQCFS
RYPDHMKQHDFFKSAMPEGYVQERTIFFKDDGNYKTRAEVKFEGDTLVNRIELKGIDFKED
GNILGHKLEYNYNSHNVYIMADKQKNGIKVNFKIRHNIEDGSVQLADHYQQNTPIGDGPVLL
PDNHYLSTQSKLSKDPNEKRDHMVLLEFVTAAGITLGMDELYKGSGIGTGFPFDPHYVEVL
GERMHYVDVGPRDGTPVLFLHGNPTSSYVWRNIIPHVAPTHRCIAPDLIGMGKSDKPDLGY
FFDDHVRFMDAFIEALGLEEVVLVIHDWGSALGFHWAKRNPERVKGIAFMEFIRPIPTWDE
WAP

Calmodulin; M13; cpHaloΔ; linker; mEGFP

>Caprola2$_{ON}$
(M)SKRDAREMFQAFRTGSDQLTEEQIAEFKEAFSLFDKDGDGTITTKELGTVMRSLGQNPT
EAELQDMINEVDADGDGTIDFPEFLTMMARKMKDTDSEEEIREAFRVFDKDGNGYISAAEL
RHVMTNLGEKLTDEEVDEMIREADIDGDGQVNYEEFVVMMTAKEFPPPPPPPPPPPPPPPP
PPPPPPPPPPPPPPPPGGSRVDSSRRKFNKTAHALRAIGRLSSLEGGSSKRDAREMFQAFR
TTDVGRKLIIDQNVFIEGTLPMGVVRPLTEVEMDHYREPFLNPVDREPLWRFPNELPIAGEP
ANIVALVEEYMDWLHQSPVPKLLFWGTPGVLIPPAEAARLAKSLPNCKAVDIGPGLNLLQED
NPDLIGSEIARWLSTLEIGSGMVSKGEELFTGVVPILVELDGDVNGHKFSVSGEGEGDATYG
KLTLKFICTTGKLPVPWPTLVTTLTYGVQCFSRYPDHMKQHDFFKSAMPEGYVQERTIFFKD
DGNYKTRAEVKFEGDTLVNRIELKGIDFKEDGNILGHKLEYNYNSHNVYIMADKQKNGIKVN
FKIRHNIEDGSVQLADHYQQNTPIGDGPVLLPDNHYLSTQSKLSKDPNEKRDHMVLLEFVTA
AGITLGMDELYKGSGIGTGFPFDPHYVEVLGERMHYVDVGPRDGTPVLFLHGNPTSSYVW
RNIIPHVAPTHRCIAPDLIGMGKSDKPDLGYFFDDHVRFMDAFIEALGLEEVVLVIHDWGSAL
GFHWAKRNPERVKGIAFMEFIRPIPTWDEWAP

Hpep; Calmodulin; M13; cpHalo; linker; mEGFP

>postsyn-Caprola$2_5$
(M)LEVKEASPTSIQISWVLHLRHVRYYRITYGETGGNSPVQEFTVPGSKSTATISGLKPGVD
YTITVYAVTIFSAYRSAWPPISINYRTGTDYKDDDDKGYPYDVPDYAGSGGSGGSGSKRDA
REMFQAFRTGSDQLTEEQIAEFKEAFSLFDKDGDGTITTKELGTVMRSLGQNPTEAELQDM
INEVDADGDGTIDFPEFLTMMARKMKDTDSEEEIREAFRVFDKDGNGYISAAELRHVMTNL
GEKLTDEEVDEMIREADIDGDGQVNYEEFVVMMTAKEFPPPPPPPPPPPPPPPPPPPPPPP
PPPPPPPGGSRVDSSRRKFNKTAHALRAIGRLSSLEGGSDVGRKLIIDQNVFIEGTLPMGVV
RPLTEVEMDHYREPFLNPVDREPLWRFPNELPIAGEPANIVALVEEYMDWLHQSPVPKLLF
WGTPGVLIPPAEAARLAKSLPNCKAVDIGPGLNLLQEDNPDLIGSEIARWLSTLEIGSGMVS
KGEELFTGVVPILVELDGDVNGHKFSVRGEGEGDATNGKLTLKFICTTGKLPVPWPTLVTTL
GYGVACFARYPDHMKQHDFFKSAMPEGYVQERTISFKDDGTYKTRAEVKFEGDTLVNRIVL
KGIDFKEDGNILGHKLEYNFNSHKVYITADKQKNGIKANFKTRHNVEDGGVQLADHYQQNT
PIGDGPVLLPDNHYLSHQSKLSKDPNEKRDHMVLKERVTAAGITHDMDELYKGSGIGTGFP
FDPHYVEVLGERMHYVDVGPRDGTPVLFLHGNPTSSYVWRNIIPHVAPTHRCIAPDLIGMG
KSDKPDLGYFFDDHVRFMDAFIEALGLEEVVLVIHDWGSALGFHWAKRNPERVKGIAFMEFI
RPIPTWDEWAPGSGGSGGSGFQCRICMRNFSDRSNLSRHIRTHTGEKPFACDICGRKFAIS
SNLNSHTKIHTGSQKPFQCRICMRNFSRSDNLARHIRTHTGEKPFACDICGRKFATSGNLTR
HTKIHLRGSQLSIVDAPEQREGASQVSVSVTFEDVAVLFTRDEWKKLDLSQRSLYREVMLE
NYSNLASMA

PSD95.FingR; FLAG; HA-tag; Hpep; Calmodulin; M13; cpHaloΔ; linker; mGreenLantern; ZF(CCR5-L); KRAB(A)

<u>M13 sequences</u>

| | |
|---|---|
| CaprolaX$_1$ | RVDSSRRKWNKTGHALRAIGRLSSLE |
| CaprolaX$_2$ | RVDSSRRKWNKTGHAVRAIGRLSSLE |
| CaprolaX$_3$ | RVDSSRRKFNKAGHALRAIGRLSSLE |
| CaprolaX$_4$ | RVDSSRRKFNKTGHAVRAIGRLSSLE |
| CaprolaX$_5$ | RVDSSRRKFNKTAHALRAIGRLSSLE |
| CaprolaX$_6$ | RVDSSRRKFNKTGHALRAIGRLSSLE |
| CaprolaX$_7$ | RVDSSRRKLNKTGHALRAIGRLSSLE |
| CaprolaX$_8$ | RVDSSRRKWNKTGHATRAIGRLSSLE |
| CaprolaX$_9$ | RVDSSRRKMNKTGHALRAIGRLSSLE |
| CaprolaX$_{10}$ | RVDSSRRKVNKTGHALRAIGRLSSLE |
| CaprolaX$_{12}$ | RVDSSRRKYNKTGHALRAIGRLSSLE |
| CaprolaX$_{13}$ | RVDSSRRKFNKTGKALRAIGRLSSLE |
| CaprolaX$_{15}$ | RVDSSRRKFNKDGHALRAIGRLSSLE |

<u>Localization sequences</u>

>Nuclear export sequence (NES, N-terminally)
(M)LQNELALKLAGLDINKT

<u>Purification sequences</u>

>pET-51b(+) N-term (Strep-tag) (Caprola1)
MASWSHPQFEKGADDDDKVPHGGS
Strep-tag; linker

>pET-51b(+) C-term (His-tag) (Caprola1)
APGFSSISAHHHHHHHHHH
Linker; His-tag

>pET-51b(+) N-term (His-tag) (Caprola2 and Caprola3)
MHHHHHHHHHHGSG
His-tag; linker
